# High-Content Screening Identifies Dithiocarbamates As A Class Of Chemicals That Disrupts TDP-43 Proteostasis

**DOI:** 10.64898/2026.08.14.741835

**Authors:** Giulia Fragola, Ryan D. Weeks, Justin Wolter, Audra F. Bryan, Katheryn N. Kapfer, Xu Tian, Julie C. Necarsulmer, Baggio A. Evangelista, Vanya Bhat, Omeed K. Arooji, Adriana S. Beltran, Tara A. Brennan, Matthew J. Niederhuber, Austin Hepperla, Leonard B. Collins, Taufika Islam Williams, Ashley J. Ezzell, Antonio Planchart, Todd J. Cohen

## Abstract

Transactive response DNA-binding protein 43 (TDP-43) aggregation and loss of function are hallmark features of amyotrophic lateral sclerosis (ALS) and frontotemporal dementia (FTD) among other neurodegenerative diseases. Despite epidemiological evidence linking environmental exposures to neurodegeneration, few toxicants have been directly associated with neurodegeneration. Here, we performed a high-content imaging screen, using a library of over a thousand chemical compounds that are considered high risk for human exposure and identified 21 toxicants that drive TDP-43 aggregation. Among the top chemical hits, five belonged to the dithiocarbamate (DTC) class of thiol-reactive compounds including the agricultural pesticides thiram and ziram. Thiram directly promoted TDP-43 cysteine oxidation and intermolecular crosslinking, whereas ziram induced TDP-43 aggregation via zinc imbalance and enhanced oxidative stress, suggesting DTCs disrupt redox homeostasis. In primary neurons and human iPSC-derived neurons, DTCs led to TDP-43 aggregation and prominent splicing defects consistent with loss of TDP-43 function. In exposed zebrafish, DTCs impaired TDP-43 function and triggered widespread transcriptional changes reflected by perturbed stress response and metabolic signatures. By combining TDP-43 loss of function mutations with chemical exposures, we observed accelerated TDP-43 loss of function and chemical-induced aggregation, supporting a multiple hit mechanism driving TDP-43 dysfunction. Together, these findings identify DTCs, particularly those used as agricultural pesticides, as dominant modifiers of TDP-43 proteostasis and identify redox imbalance and zinc homeostasis as a central molecular mechanism linking toxicant exposure to TDP-43 proteinopathy.

## Introduction

The pathological mislocalization and aggregation of Transactive Response DNA-binding protein 43 kDa (TDP-43) has emerged as a defining hallmark shared across a wide spectrum of neurodegenerative disorders [1, 2]. Initially identified as a pathological hallmark in frontotemporal dementia (FTD) and amyotrophic lateral sclerosis (ALS) [3], TDP-43 pathology is now recognized in multiple degenerative conditions, including Alzheimer’s disease (AD)[4], chronic traumatic encephalopathy (CTE)[5], inclusion body myositis (IBM) and Limbic-predominant Age-related TDP-43 Encephalopathy (LATE)[6], that are collectively referred to as TDP-43 proteinopathies [2].

TDP-43 dysfunction is characterized by the loss of its normal diffuse nuclear localization, followed by the accumulation of cytoplasmic and intranuclear TDP-43–positive inclusions. Studies indicate that nuclear depletion and loss of TDP-43 function precede overt aggregation [3, 7–9]. These observations suggest that early nuclear TDP-43 dysfunction is a key pathogenic event driving downstream cellular death. The consequences of TDP-43 loss of function are extensive, given its central role as a highly conserved, ubiquitously expressed RNA-binding protein that predominantly resides in the nucleus while dynamically shuttling to the cytoplasm [10]. TDP-43 participates in multiple aspects of RNA metabolism [11] and is estimated to regulate more than 6,000 transcripts in the brain [12], including autoregulation of its own mRNA [13], as well as transcripts critical for neuronal function and disease pathology, such as *STMN2* [14–16]*, SORT1* [12] and *KCNQ2* [17].

The propensity of TDP-43 to undergo phase separation and pathological aggregation is dictated by the structural properties of its protein domains. TDP-43 contains an N-terminal domain (NTD) [18], two RNA-recognition motifs (RRMs) required for RNA binding [19], and a glycine-rich C-terminal domain (CTD) [20]. Conformational changes within the CTD strongly influence aggregation behavior [10], consistent with the observation that most disease-linked mutations and post-translational modifications map to this region [21]. We previously identified cysteine oxidation and disulfide cross-linking as an early event that promotes TDP-43 aggregation [22], suggesting redox-sensitive cysteine residues can act as reversible switches for stress signaling. Consistent with a stress-dependent mechanism, oxidative stressors such as sodium arsenite trigger TDP-43 cysteine oxidation, intra-and intermolecular disulfide bond formation, and subsequent aggregation. These findings implicate oxidative stress and thiol-reactive TDP-43 cysteines as putative sensors by which environmental chemicals could directly perturb TDP-43 function in disease.

The contribution of environmental exposures to the development of FTD/ALS is increasingly recognized [23]. For example, exposure to β-methylamino-L-alanine (BMAA), a non-proteinogenic amino acid produced by cyanobacteria, has been linked to a highly penetrant form of ALS with parkinsonism and dementia among Chamorro populations consuming BMAA-contaminated diets [24]. Neuropathological examination of affected individuals revealed extensive TDP-43–positive neuronal inclusions and dystrophic neurites accompanied by loss of normal nuclear TDP-43 localization [25, 26]. Even beyond BMAA, studies have shown that a variety of metals and industrial chemicals, including arsenite, lead, mercury, zinc, and dioxins, can influence TDP-43 expression, solubility, or aggregation state [27–31]. Moreover, epidemiological reports have shown that persistent environmental pollutants are associated with an increased risk of developing ALS [32]. Despite these observations, a systematic identification of environmental compounds that promote TDP-43 pathology, particularly high-risk toxicants to which humans are routinely exposed, remains largely unexplored.

To address this gap, we performed a high-content imaging screen to identify toxicants capable of inducing TDP-43 aggregation. We screened a curated library of 1,052 environmental compounds classified as at-risk by the U.S. Environmental Protection Agency Toxicity Forecaster (ToxCast) program [33]. Primary candidates were subsequently validated across multiple cellular models and neuronal assays, enabling the identification of compounds that induce gain and loss of function TDP-43 toxicity. Among the eight chemicals that consistently promoted TDP-43 aggregation, five belonged to the dithiocarbamate (DTC) class of thiol-reactive compounds: thiram, ziram, sodium dimethyldithiocarbamate (SDTC), disulfiram, and dazomet. The agricultural pesticides thiram and ziram depleted soluble TDP-43 and led to TDP-43 aggregation via cysteine oxidation, leading to impaired TDP-43–dependent splicing function in mouse and human induced pluripotent stem cell (iPSC)-derived cortical neurons. In exposed adult zebrafish, DTCs elicited toxicant-specific phenotypes as well as common transcriptional signatures that may link this family of chemicals to TDP-43 dysfunction.

## Results

### A high throughput screen identifies chemical inducers of TDP-43 aggregation

To identify toxicants linked to TDP-43, we acquired a library of 1,052 prioritized chemical compounds from the EPA (ToxCast phase II) [33] and tested their ability to induce TDP-43 aggregation in a well characterized TDP-43 reporter line in which HEK-293A cells were engineered to stably express a doxycycline (dox) inducible GFP tagged TDP-43 containing a mutated nuclear localization signal (iGFP-TDP43-dNLS) [34] (Fig. 1 and Supplementary Fig. 1). In this cellular model, TDP-43 is predominantly expressed in the cytoplasm upon doxycycline addition. Cells were cultured with doxycycline for 24 hours, followed by a 6-hour exposure to three different concentrations (4-20-100 µM) of individual ToxCast compounds in a triplicate well format (Fig. 1b). Arsenite was added as a positive control for TDP-43 aggregation. After exposure, cells were simultaneously fixed and permeabilized (1%Triton) to remove soluble TDP-43 while retaining insoluble aggregated TDP-43 and to a lesser extent nuclear RNA-bound TDP-43 [34] (Fig. 1b). We used a high content imaging system (IN Cell Analyzer 2000) to examine GFP-positive TDP-43 from an average of 1,125 cells/well. The total number of GFP-positive cells per well and the percentage of cells with TDP-43-GFP aggregates were calculated using a customized CellProfiler pipeline (Fig. 1 and Supplementary Fig. 1). A Z-factor of 0.78 was observed between control (DMSO) and arsenite (100 µM) treated wells, supporting the overall assay quality and dynamic range of the screen (Supplementary Fig. 1b).

**Figure 1.**
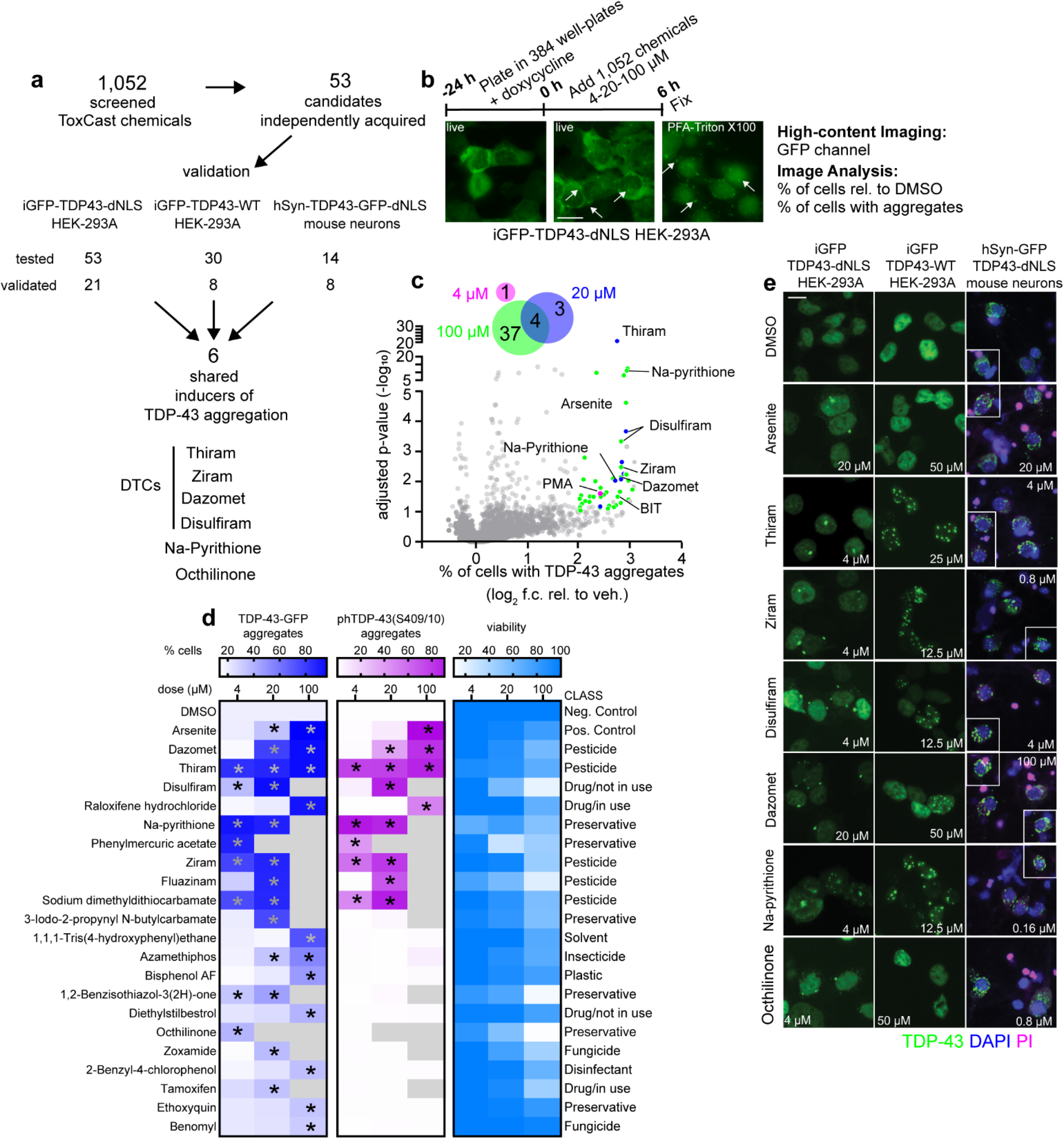
Identification of at-risk environmental chemicals that promote TDP-43 aggregation. **a** Screening results summary. **b** Screening protocol. Scale bar = 25 µm**. c** Scatter plot showing the identification of 45 candidate environmental chemicals promoting TDP-43 pathology without overt toxicity. Venn diagram shows number of hits identified at each concentration. Scatter plot shows pooled data of 3 chemical concentrations (4-20-100 µM), each dot is the mean of 3 replicates for each concentration. X-axis is the log_2_ fold change of the % cells with TDP-43 aggregates, y-axis is-log_10_ adjusted p-values (two-tailed t-test followed by a Benjamini-Hochberg’s false discovery rate adjustment). Green = 41 hits at 100 µM, blue = 7 hits at 20 µM, magenta = 1 hit at 4 µM, grey = chemicals not identified as hits. 4 chemicals were identified as hits at both 20 µM and 100 µM concentrations. ctr. = control, f.c. = fold change, veh. = vehicle = DMSO, BIT = 2-Benzisothiazol-3(2H)-one, PMA = Phenylmercuric acetate. **d** Identification of 21 chemical inducers of TDP-43 aggregation by validation of independently acquired chemical candidates in iGFP-TDP43-dNLS HEK-293A cells. Heatmap showing the percentage of cells with TDP-43-GFP aggregates (blue), phTDP-43(S409/10) aggregates (magenta) and the number of cells/well as percentage relative to vehicle wells (light blue), for DMSO, each of the 21 chemicals and arsenite at 3 different concentrations (4-20-100 µM) after 6 hours of treatment. Shown chemicals induce a significant increase in TDP-43-GFP aggregates with less than 50% viability loss in at least one dose. Hits are ranked based on the degree of aggregation induced at statistically significant doses. Two-way ANOVA followed by Dunnett’s multiple comparison test. **e** Representative images of iGFP-TDP43-dNLS HEK-293A cells (left), iGFP-TDP43-WT HEK-293A cells (center) and hSyn-TDP43-dNLS-GFP infected neurons (right) treated with DMSO or the indicated concentration of a common set of chemicals inducing TDP43 aggregation in all 3 in vitro systems. Arsenite is also shown. Propidium iodide (PI, magenta) was added to hSyn-TDP43-dNLS-GFP neurons before fixation to quantify viability. White squares show neurons cropped from the same field and added. Scale bar = 25 µm. Further statistical information is available in the Statistical Data File. Extended data are available in the Extended Data File.

Analysis of the pooled data identified a total of 55 exposure conditions in which an individual chemical, at any one of three doses, induced a significant increase in the percentage of cells containing TDP-43 aggregates (-log_2_ fold changes >2,-log_10_ adjusted p-value >1) (Fig. 1 a, c). As post-screen confirmation, we manually assessed the presence of TDP-43 aggregates in response to each of the chemicals of interest and excluded two conditions where enhanced GFP fluorescence likely occurred in response to cell death (Supplementary File 1a).

The post-screen processing led to the identification of 45 unique candidate chemicals promoting TDP-43 aggregation (Fig. 1 a, c and Supplementary File 1a). In addition to those that met the p-value threshold, we included an additional 18 chemicals that showed aggregation trends that were not considered significant (Supplementary File 1b). Among the chemicals of interest, we were able to independently acquire new lots of 53 chemicals that were used to validate their ability to drive TDP-43 aggregation (Supplementary File 1b). In a follow-up analysis with all 53 individual chemicals, iGFP-TDP43-dNLS reporter cells were re-exposed for 6 hours at 4-20-100 µM (Fig. 1, Supplementary Fig. 2, 3, 4) following the same screening pipeline (Fig. 1 d, e). This led to the validation of 21 chemicals that induced a significant increase in TDP-43 aggregation without causing more than 50% cell loss (Fig. 1 d, e and Supplementary File 1c). Robust induction of TDP-43 aggregation was associated with hyperphosphorylation at S409/10 (phTDP-43(S409/10)), a hallmark of TDP-43 pathology, as shown for dazomet, thiram, disulfiram, raloxifene hydrochloride, Na-pyrithione, phenylmercuric acetate, ziram, flunazinam and SDTC (Fig. 1d, Supplementary Fig. 2).

To determine the effect on wild-type (WT) nuclear localized TDP-43, HEK-293A cells expressing an inducible GFP-tagged WT TDP-43 (iGFP-TDP43-WT) [34] were similarly exposed to a subset of 30 chemicals of interest. Since nuclear TDP-43 has a higher threshold for aggregation compared to cytoplasmic TDP-43, we increased the dosing paradigm to 12.5-50 µM for iGFP-TDP43-WT expressing cells (Fig. 1e and Supplementary Fig. 3), which avoids the occasional toxicity observed at 100 µM while providing maximum sensitivity to detect subtle changes in aggregation. Among the 30 chemicals evaluated in iGFP-TDP43-WT cells, 8 showed a significant increase in nuclear TDP-43 puncta formation from at least one of the concentrations tested: thiram, ziram, Na-pyrithione, disulfiram, dazomet, octhilinone, gentian violet, and fluazinam (Fig. 1e and Supplementary Fig. 3 and Supplementary File 1c).

To confirm the impact of any putative pro-aggregation chemicals in neurons, we employed mouse primary cortical neurons that were virally transduced with dNLS-TDP-43-GFP driven by a neuron-specific promoter (human Synapsin 1, hSyn) (Fig. 1e, Supplementary Fig. 4). Each chemical was evaluated at 5 different concentrations ranging from 0.16 – 100 µM including propidium iodide (PI) to monitor neuronal viability. While control DMSO-treated cells showed diffusely expressed cytoplasmic dNLS-TDP-43-GFP, 8 of the chemicals tested led to robust TDP-43 aggregation in neurons, without causing overt toxicity: thiram, ziram, Na-pyrithione, disulfiram, dazomet, octhilinone, SDTC and,2-benzisothiazol-3(2H)-one or (BIT) (Fig. 1e, Supplementary Fig. 4 and Supplementary File 1c).

The DTCs emerged from the above results as the most common chemical superfamily capable of driving TDP-43 aggregation. Two of its family members, thiram (CAS 137-26-8) and ziram (CAS 137-30-4) are high production agricultural pesticides widely used in the cultivation of apples, grapes, tobacco, soybeans, and wheat (U.S. Geological Survey). Dazomet (CAS 533-74-4) is a fumigant and pesticide mainly used for soil sterilization. Disulfiram (CAS 97-77-8) has been used to treat alcohol addiction (Antabuse) and has been recently repurposed for cancer treatment [35, 36]. Sodium dimethyldithiocarbamate (SDDC) (CAS 128-04-1) is a high production chemical used as a pesticide and chelating agent to treat wastewater.

### DTCs induce cysteine-dependent TDP-43 aggregation via distinct mechanisms

Our prior study identified TDP-43 cysteine-dependent redox regulation in response to stressors [22]. Among the most significant chemicals identified in vitro, five are known to have thiol reactive properties (thiram, ziram, SDTC, disulfiram, dazomet and BIT) [37–42]. To test direct toxicant reactivity with TDP-43 cysteines, we exposed recombinant TDP-43 (rTDP-43) to each individual chemical (thiram, ziram, disulfiram, SDTC, Na-pyrithione and BIT) in vitro for 1 h, followed by labeling with 5-iodoacetamidofluorescein (5-IAF), which fluorescently labels free thiol groups and therefore acts as a readout for cysteine reactivity (Fig. 2 a-e, Supplementary Fig. 5). In the absence of chemical exposure, rTDP-43 was primarily detected as monomer (M; ∼43 kDa) with subtle accumulation of multimeric species (trimer ∼130 kDa, tetramer ∼170 kDa), including minimal high-molecular-weight species (HMW; >250 kDa), the latter of which resolved to monomeric and oligomeric forms under reducing conditions (Fig. 2 a-e, Supplementary Fig. 5).

**Figure 2.**
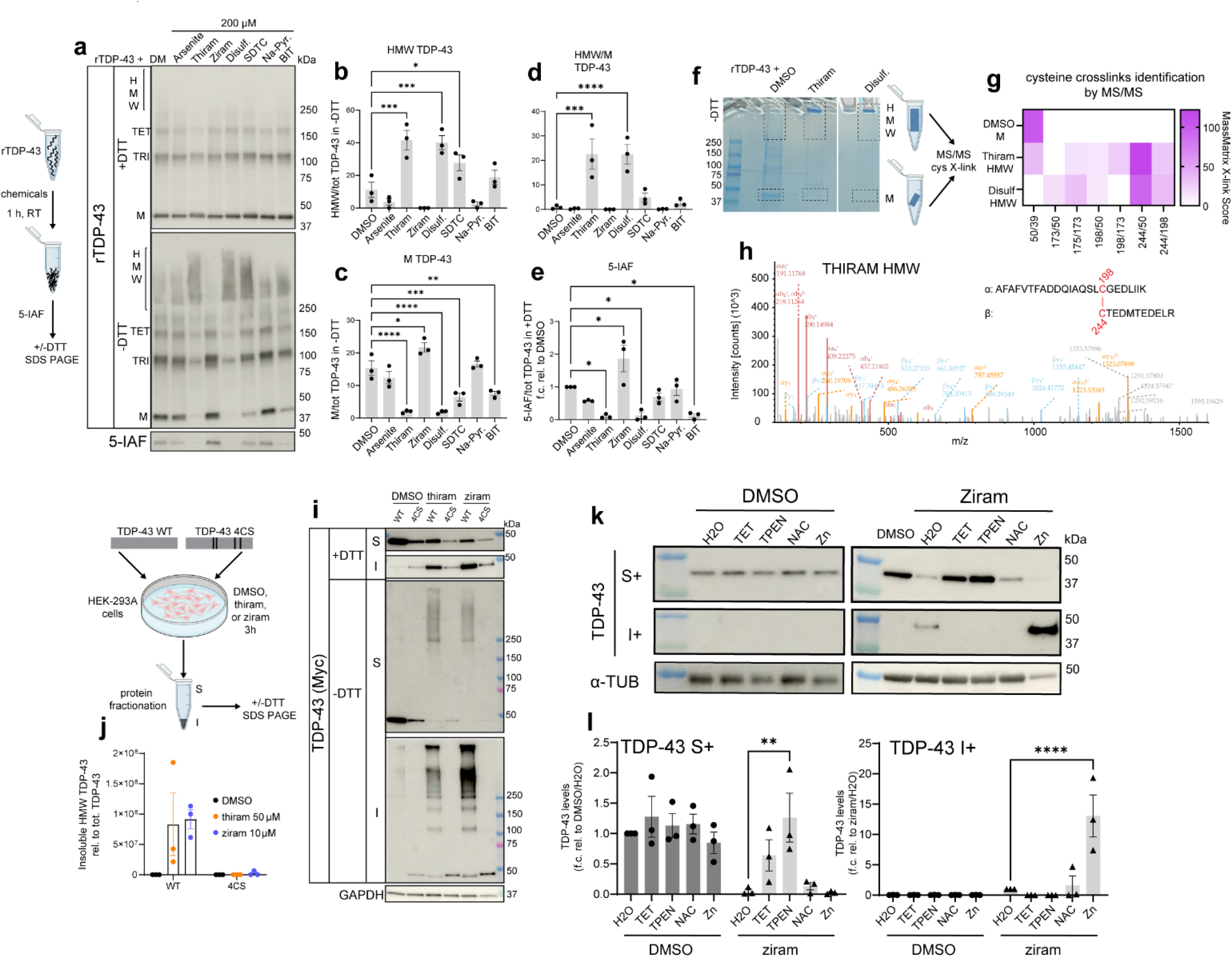
DTCs induce cysteine-dependent TDP-43 aggregation via distinct mechanisms. **a** Representative Western blot images of rTDP-43 after 1 hour incubation with DMSO or 200 uM of the indicated chemicals followed by addition of 80 µM of 5-Iodoacetamidofluorescein (5-AIF). Monomeric (M; 43 kDa), high molecular weight (HMW; >250 kDa), trimeric (TRI, ∼130 kDa), tetrameric (TET; ∼170 kDa). 5-IAF levels are also shown. Bar-charts show quantification of HMW (**b**) and M (**c**) TDP-43 species normalized to total amounts of TDP-43 in-DTT blots, HMW/M TDP-43 ratio (**d**) and amounts of 5-IAF signal normalized to M TDP-43 in +DTT blots (**e**). Bars represent mean ± SEM; dots represent n=3 biological replicates. One-way ANOVA followed by Dunnet’s multiple comparison test. **f** Comassie gel of rTDP-43 treated with DMSO or 20 uM of thiram or disulfiram for 1 hour, followed by non-reducing SDS-PAGE. TDP-43 HMW and monomer were isolated for MS/MS to identify cysteines crosslinks. **g** Heatmap showing X-link scores of cysteines crosslinks identified by MassMatrix in the indicated samples. **h** Representative peptide MS spectra of C198-C244 crosslink identified in thiram treated rTDP-43. **i** Representative Western blot images of HEK-293A cells transfected with Myc-tagged TDP-43 WT or C173/175/189/277S (4CS) and treated with DMSO, 50 µM of thiram or 10 µM of ziram for 3 hours. Lysates were fractionated in RIPA soluble (S) and insoluble (I) fractions and analyzed by Western blot in the presence or absence of DTT. **j** Quantification of insoluble HMW aggregates normalized to total amounts of Myc-TDP-43 in each sample (sum of S+ and I+ signal). Bars represent mean ± SEM; dots represent n=3 biological replicates. One-way ANOVA followed by Dunnet’s multiple comparison test. **k** Representative Western blot images of mouse primary neurons pretreated with 120 µM triethylenetetramine hydrochloride (TET), 25 µM of N,N,N′,N′-tetrakis(2-pyridylmethyl)ethylenediamine (TPEN), 100 µM of N-acetylcysteine (NAC), or 100 µM of zinc (Zn). Neurons were then treated with DMSO (left) or 7.5 µM of ziram (right) for 3 hours. **l** Quantification of TDP-43 soluble and insoluble amounts normalized to α-TUBULIN and expressed as fold change relative to control. Bars represent mean ± SEM; dots represent n=3 biological replicates. Two-way ANOVA followed by Dunnet’s multiple comparison test. *p < 0.05, **p < 0.01, ***p < 0.001, ****p < 0.0001. Only significant p values are shown. Further statistical information is available in the Statistical Data File. Extended data are available in the Extended Data File.

Thiram and disulfiram induced a marked increase in the HMW/monomer ratio (Fig. 2a, d), driven by accumulation of HMW species (Fig. 2a, b) and a concomitant decrease in monomeric TDP-43 (Fig. 2a, c). These HMW species were reduced to monomeric TDP-43 in the presence of the reducing agent dithiothreitol (DTT) (Fig. 2a), indicating reversible cysteine crosslinking. Consistently, both thiram and disulfiram significantly decreased free cysteine availability (Fig. 2a, e), supporting direct toxicant reactivity with TDP-43 cysteines and the formation of intermolecular cysteine-mediated crosslinking and aggregation. SDTC produced more modest effects on HMW and monomer levels without altering the HMW/monomer ratio or cysteine availability (Fig. 2 a-e), suggesting a weaker or transient cysteine interaction.

In contrast, ziram did not induce HMW accumulation in this cell-free system but instead modestly increased monomer levels and accessible cysteine residues, consistent with a potential conformational change that enhances free thiol exposure (Fig. 2 a-e, Supplementary Fig. 5). BIT decreased monomer levels and free cysteine availability without affecting the HMW/monomer ratio, suggesting preferential intramolecular crosslinking. As expected, Na-pyrithione, had no detectable effect on TDP-43 multimer formation (Fig. 2a-e, Supplementary Fig. 5).

To identify the relevant cysteines involved in crosslinks, we performed MS/MS analysis on monomeric (∼43 kDa) and HMW (>250 kDa) TDP-43 species following treatment with control (DMSO), thiram, or disulfiram (Fig. 2 f-h). Consistent with the in vitro immunoblotting analysis, DMSO-treated samples were mostly monomeric TDP-43, whereas thiram and disulfiram induced a near-complete redistribution of TDP-43 into HMW species by Coomassie staining (Fig. 2f). Control (DMSO) samples exhibited minimal cysteine crosslinking (primarily C39–C50), while thiram and disulfiram treated samples showed a marked increase in the number and crosslink branches detected, with C50–C244 and C198–C244 predominating (Fig. 2g, h). Thus, thiram and disulfiram exposures promote intermolecular cysteine-mediated TDP-43 crosslinking, consistent with their ability to drive TDP-43 aggregation.

To determine whether chemical exposure similarly involves cysteines in a cellular microenvironment, HEK-293A cells overexpressing either wild-type (WT) TDP-43 or a cysteine-deficient TDP-43 mutant (4CS; C173/175/198/244A) [22] were exposed to thiram (50 µM) or ziram (10 µM) for 3 h, and RIPA soluble or insoluble cell lysates were analyzed by immunoblotting (Fig. 2i-j). Both thiram and ziram induced soluble and insoluble HMW TDP-43 species in WT-expressing cells, whereas HMW aggregate formation was completely abolished in cells expressing the 4CS mutant (Fig. 2i-j). Notably, ziram induced cysteine-dependent aggregation in cells (Fig. 2i-j) but not in the cell-free system (Fig. 2a-e), indicating that ziram is unlikely to directly chemically modify TDP-43, which is more consistent with ziram causing stress via an indirect mechanism. These data suggest that DTC-induced TDP-43 aggregation is cysteine-dependent and can occur via either direct chemical modification of TDP-43 (thiram, disulfiram) or via indirect cellular mechanisms (ziram).

### Thiram and ziram induce TDP-43 loss of function in mouse and human neurons

To determine whether DTCs impact endogenous TDP-43 in neurons, we exposed wild type mouse cortical neurons to increasing doses of thiram, ziram and disulfiram for 3 hours and examined TDP-43 solubility and phosphorylation at Ser409/410 by standard immunoblotting of soluble and insoluble fractions (Supplementary Fig. 6a, b). Formation of HMW TDP-43 aggregates was assed using non-reducing immunoblotting. All 3 chemicals induced a dose dependent shift of TDP-43 from the soluble to the insoluble fraction with concomitant hyperphosphorylation and formation of crosslinked HMW aggregates (Supplementary Fig. 6 a, b). We focused our follow-up mechanistic analysis specifically on thiram and ziram given their widespread use as agricultural pesticides and their prior environmental link to neurodegeneration [37, 43–45].

Ziram is a cell-permeant complex comprised of zinc (Zn) with two dimethyldithiocarbamate ligands [46]. Because of its metal-binding capabilities, ziram chelates and transports copper and zinc, leading to intracellular metal imbalance and consequent oxidative injury [46]. To test if ziram indirectly promotes TDP-43 aggregation via increased intracellular copper or zinc, mouse neurons were pretreated with the cell permeant copper chelator (triethylenetetramine hydrochloride-TET) or zinc chelator (N,N,N′,N′-tetrakis(2-pyridylmethyl)ethylenediamine - TPEN) prior to ziram exposure. We included a pretreatment with the antioxidant N-acetyl-L-cysteine (NAC) as a comparison. In addition, to further assess if zinc exacerbates TDP-43 aggregation, we included a separate condition in which zinc itself was supplemented in the media prior to ziram exposure (Fig. 2 k, l). Pretreatment with TPEN led to a complete rescue of ziram-induced TDP-43 aggregation, as indicated by restored soluble TDP-43 and elimination of the insoluble TDP-43 pool. Conversely, pretreatment with zinc potentiated ziram’s effect on TDP-43 aggregation. TET and NAC showed more intermediate phenotypes including a partial rebalancing of TDP-43 solubility, as soluble TDP-43 levels were partly restored, though this effect was not statistically significant. Importantly, none of the chelator or metal exposures had any effect on TDP-43 solubility in the absence of ziram (Fig. 2 k, l). These data suggest that zinc metabolism is downstream of ziram-induced TDP-43 aggregation.

ALS/FTD is considered multifactorial in which susceptibility is dictated by the interaction of environmental and genetic factors [23, 47]. To explore possible gene-environment interactions, we tested whether a genetically encoded TDP-43 RNA-binding deficient mutation (K145Q), previously characterized by our group [48], would confer increased sensitivity to chemical exposure. We therefore cultured mouse cortical neurons from TDP-43^K145Q^ mice and their respective wild-type littermate controls and exposed them to thiram and ziram (25-50 µM for thiram, 3.3-7.5 µM for ziram) (Fig. 3, Supplementary Fig. 7). Ziram did not significantly impact cell viability at either concentration, whereas thiram caused a ∼ 30% decrease in viability at the highest dose (Fig. 3a). At baseline, TDP-43^K145Q^ neurons exhibited higher levels of soluble TDP-43 than WT, confirming that the TDP-43^K145Q^ mutation perturbs RNA binding and prevents autoregulation of its own *T*ardbp transcript (Fig. 3b, Supplementary Fig. 7b). To characterize the impact on neuronal health, we monitored heme oxygenase-1 (HO-1) levels as a surrogate measure of oxidative stress. At baseline, levels of HO-1 were comparable between WT and TDP-43^K145Q^ neurons (Fig. 3b, Supplementary Fig. 7f). Upon treatment with 50 µM thiram, a robust and significant HO-1 increase was observed in TDP-43^K145Q^ neurons, suggesting a lower threshold for thiram-induced dysfunction in the setting of a partially impaired mutant TDP-43 (Fig. 3b, c, Supplementary Fig. 7e). Similarly, ziram did not induce HO-1 levels in WT neurons but showed a modest increase in TDP-43^K145Q^ mutant neurons that was not statistically significant (Fig. 3b, c, Supplementary Fig. 7e).

**Figure 3.**
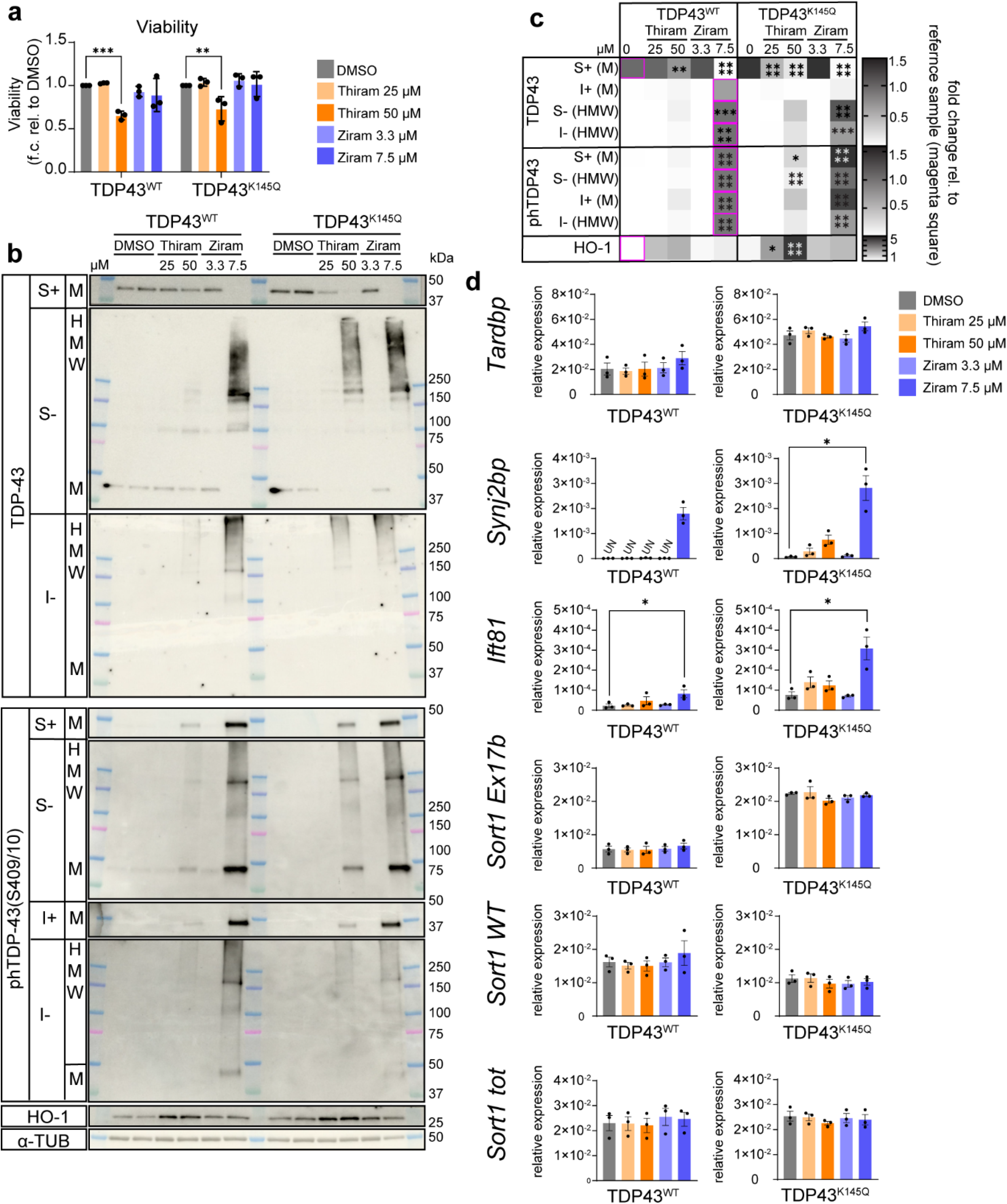
Thiram and ziram induce TDP-43 loss of function in primary mouse neurons. **a** Cell viability measured by CellTiter-Blue assay performed on WT and TDP43^K145Q^ mouse cortical neurons after 3 hours of exposure of to the different treatments listed. Bars represent mean ± SEM; dots represent n=3 biological replicates. Two-way ANOVA (genotype x treatment) followed by Dunnet’s multiple comparison test. **b** Representative Western blot images of WT and TDP43^K145Q^ mouse cortical neurons treated with DMSO or 2 doses of thiram (25-50 µM) or ziram (3.3-7.5 µM) for 3 hours. Soluble (S) and insoluble (I) protein fractions were run in presence (+) or absence (-) of DTT. Membranes were blot with TDP-43, phTDP-43(S409/10), HO-1 and α-TUBULIN (loading control) antibodies. Monomeric (M) and high molecular weight (HMW) TDP-43 species are indicated on the left. TDP-43 I+ representative images are not shown for lack of consistency across biological replicates. **c** Quantification of monomeric (M) or aggregated (HMW) TDP-43, phTDP-43(S409/10) and HO-1 protein levels in soluble and insoluble protein fractions normalized to α-TUBULIN. Heatmap shows mean of fold changes in protein levels relative to WT DMSO (S+) or WT 7.5 µM ziram (S-, I+, I-). Magenta squares indicate reference samples for each row. n = 3 independent biological replicates. Two-way ANOVA (genotype x treatment) followed by Dunnet’s multiple comparison test. **d** Effect of thiram (25-50 µM) and ziram (3.3-7.5 µM) 3-hour-treatments on TDP-43 splicing function in WT and TDP43^K145Q^ mouse cortical neurons. qRT-PCR analysis showing inclusion of cryptic regions in *Synj2bp* and *Ift81* transcripts normalized to *Gapdh* housekeeping control. Bars represent mean ± SEM of 2^(-ΔCt)^ relative to *Gapdh*; dots represent n=3 biological replicates. Friedman test with Dunn’s multiple comparisons test was performed on ΔCt values. *p < 0.05, **p < 0.01, ***p < 0.001, ****p < 0.0001. Only significant p values are shown. *Synj2bp* graph for TDP-43^wt^ neurons is provided for visualization purposes only, because non-detectable (UN) levels of *Synj2bp* cryptic exon-containing transcripts in one or more replicates of the indicated conditions prevented statistical comparisons among groups. Further statistical information is available in the Statistical Data File. Extended data are available in the Extended Data File.

We next evaluated pathological TDP-43 phosphorylation and HMW aggregate formation using TDP-43 and phTDP-43(S409/10) antibodies (Fig. 3b, c, Supplementary Fig. 7c, d). Lower concentrations of ziram exposure did not alter TDP-43 solubility or phosphorylation, however, exposure to 7.5 µM ziram resulted in a near-complete loss of soluble TDP-43 (S+) and a corresponding conversion into insoluble crosslinked aggregates (S− and I−) in both WT and TDP-43^K145Q^ neurons. Loss of soluble TDP-43 was frequently, but not always, associated with the appearance of insoluble TDP-43 (Fig. 3c, Supplementary Fig. 7a, b). Ziram-induced loss of soluble TDP-43 was accompanied by phosphorylation of monomeric and HMW TDP-43 species (Fig. 3b, c, Supplementary Fig. 7c).

Exposure to thiram showed a unique TDP-43 signature dependent on WT vs. TDP-43^K145Q^ genotype. For example, in WT neurons, 50 µM thiram modestly reduced soluble TDP-43 without a clear accumulation of TDP-43 aggregates. Strikingly, however, these effects were significantly exacerbated in TDP-43^K145Q^ neurons with a marked reduction of soluble TDP-43 (25 and 50 µM thiram) and accumulation of phosphorylated TDP-43-positive species. Significant genotype differences were observed at both 25 µM and 50 µM thiram (Fig. 3b, c, Supplementary Fig. 7a, b). These data suggest additive loss of function under conditions in which TDP-43 function is impaired, either by a chemical exposure (e.g., thiram) or a genetically encoded *TARDBP* mutation. We note that the effects of 7.5 µM ziram reach saturation independent of genotype.

To assess if thiram or ziram promotes TDP-43 loss of function, we performed quantitative reverse transcription-polymerase chain reaction (qRT-PCR) analysis of known TDP-43 target genes in mice, including the autoregulated *Tardbp* transcript, *Sort1 ex17b* splicing, and cryptic exon (CE) inclusion in *Synj2bp* and *Ift81* [49, 50] (Fig. 3d, Supplementary Fig. 8a). As expected at baseline, TDP-43^K145Q^ mutant neurons show deregulated splicing for all TDP-43 targets tested, substantiating the K145Q mutant as TDP-43 loss of function (Fig. 3d, Supplementary Fig. 8a). Depending on the target gene analyzed, we noted different patterns of splicing dysregulation in response to thiram and ziram. In particular, while *Synj2bp* CE was undetectable in untreated WT neurons, treatment with 7.5 µM ziram led to detectable levels of *Synj2bp* CE in WT neurons and a significant increase over baseline expression in TDP-43^K145Q^ neurons (Fig. 3d). Treatment with 7.5 µM ziram also induced a significant induction of *Ift81* CE expression in both WT and TDP-43^K145Q^ neurons. There were also scenarios in which no splicing effects were observed. For example, thiram did not significantly affect TDP-43 splicing targets, and neither thiram nor ziram impacted *Sort1 ex17b* or *Tardbp* expression (Fig. 3d).

To determine the effect of thiram and ziram on human neurons, we performed a 3-hr exposure in WT or TDP-43^K145Q^ iPSC-derived human cortical neurons [48] (Fig. 4). Both concentrations of ziram (5 and 7.5 µM) and the highest concentration of thiram (50 µM) significantly decreased soluble TDP-43 regardless of genotype. The lower thiram exposure (25 µM) decreased TDP-43 solubility only in TDP-43^K145Q^ neurons, suggesting enhanced sensitivity (Fig. 4a, b). In general, we observed that depletion of the soluble TDP-43 pool was a much more reliable response to chemical exposure, since HMW insoluble TDP-43 aggregates showed more inter-replicate variability, which may arise from subtle differences in neuronal maturation and differentiation across biological replicates. Overall, thiram and ziram both impacted TDP-43 in mouse and human neurons, with some subtle differences in the additive effects of exposure on genotype.

**Figure 4.**
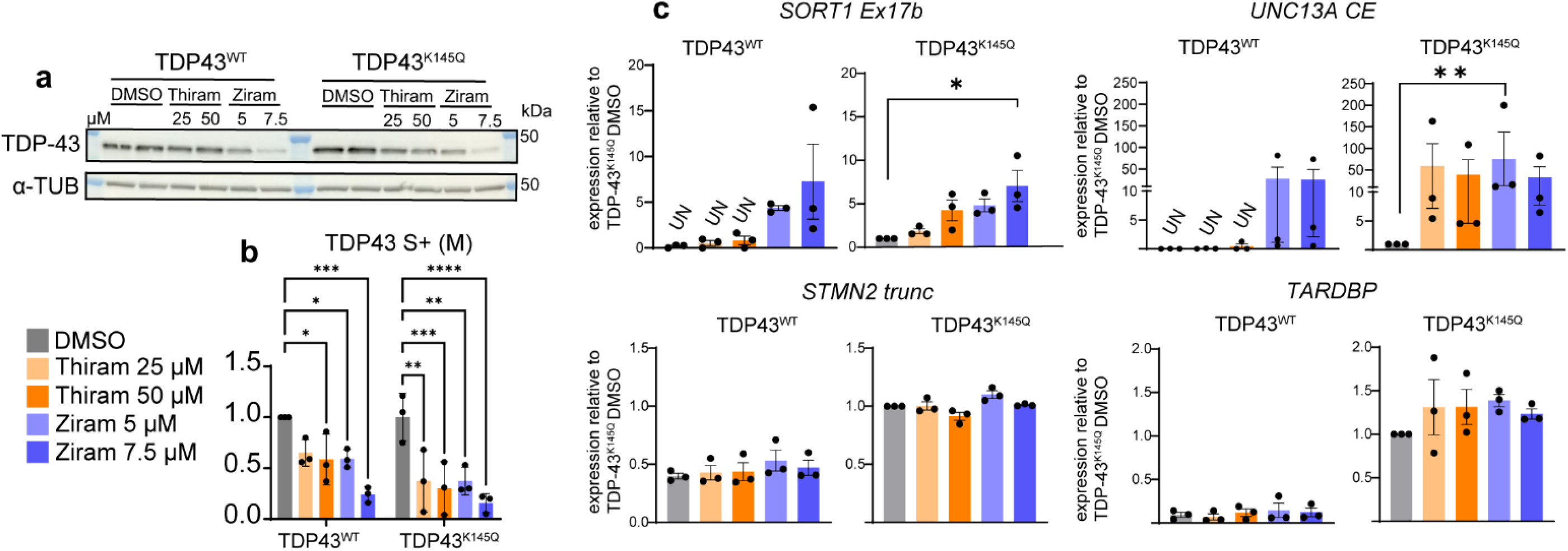
Thiram and ziram induce TDP-43 loss of function in human iPSC-derived neurons. **a** Representative Western blot images of WT and TDP43^K145Q^ hiPSC-derived cortical neurons treated with DMSO or 2 concentrations of thiram (25-50 µM) or ziram (5-7.5 µM) for 3 hours. Soluble proteins were run in presence of DTT. Membranes were blot with TDP-43 and α-TUBULIN (loading control) antibodies. **b** Quantification of soluble monomeric TDP-43 protein levels normalized to α-TUBULIN. Bars represent mean ± SEM of fold changes relative to WT DMSO condition; dots represent n=3 independent hiPSC differentiations and exposures. Only significant p values are shown. Two-way ANOVA (genotype x treatment) followed by Dunnet’s multiple comparison test. **c** qRT-PCR analysis showing amounts of the indicated transcripts in WT and TDP43^K145Q^ hiPSC-derived cortical neurons treated with thiram (25-50 µM) or ziram (5-7.5 µM) for 3 hours. Bars represent mean ± SEM of RNA levels relative to *RPLP0 a*nd normalized to TDP43^K145Q^ DMSO sample; dots represent n=3 independent iPSC differentiations and exposures. Friedman test with Dunn’s multiple comparisons test was performed on ΔCt values. *p < 0.05, **p < 0.01, ***p < 0.001, ****p < 0.0001. Only significant p values are shown. *SORT1 Ex17b and UNC13A CE* graphs for TDP-43^wt^ hiPSC-derived neurons are provided for visualization purposes only, because non-detectable (UN) levels of transcripts in one or more replicates of the indicated conditions prevented statistical comparisons among groups. Further statistical information is available in the Statistical Data File. Extended data are available in the Extended Data File.

Using iPSC-derived neurons, we next performed qRT-PCR analysis of established TDP-43-regulated human splicing events, including the FTD/ALS risk genes *STMN2*, *UNC13A* and *SORT1* [14–16, 50–52] (Fig. 4c, Supplementary Fig. 8b-9). TDP-43^K145Q^ human neurons showed a partial loss of TDP-43 function at baseline, as demonstrated by increased *TARDBP* levels (Fig. 4c, Supplementary Fig. 8b). Moreover, both *SORT1 Ex17b* and *UNC13A CE* transcripts were either undetectable or very low (cycle thresholds > 35) in WT neurons under basal conditions but always present in TDP-43^K145Q^ neurons (Fig. 4c). Treatment with ziram induced both *SORT1 Ex17b* and *UNC13A CE* expression in WT neurons (Fig. 4c). Similarly, we observed a trend toward increased *SORT1 Ex17b* and *UNC13A* CE inclusion in TDP-43^K145Q^ neurons following both thiram and ziram exposures, although only 25 µM thiram reached significance (Fig 4c). No treatment-dependent effects were detected for *TARDBP, UNC13A WT, SORT1 WT* or *STMN2* (Fig. 4c, Supplementary Fig. 9).

### DTCs alter zebrafish TDP-43 levels and cause genome-wide transcriptional perturbations

To determine whether thiram and ziram impact TDP-43 function and drive neurodegenerative phenotypes *in vivo*, we exposed adult 18-month-old zebrafish to ziram or thiram for 15 days and assessed behavioral, molecular, and transcriptional changes (Fig. 5-6). Motor function was measured by the swim tunnel assay, which revealed no detectable deficits following exposure to either chemical (Fig. 5a-c). Anxiety-like behavior was assessed by the novel tank assay that measures changes in the zebrafish natural inclination to occupy the bottom zone of a new environment (Fig. 5d-i). Ziram-treated fish displayed an anxiety-like phenotype in the novel tank assay, characterized by increased time spent in the bottom zone (zone 1), reduced time in the top zone (zone 3), decreased swimming velocity, and increased latency to enter the top zone (Fig. 5e, g, i). Thiram treatment did not significantly impact performance in the novel tank assay (Fig. 5d, f, h).

**Figure 5.**
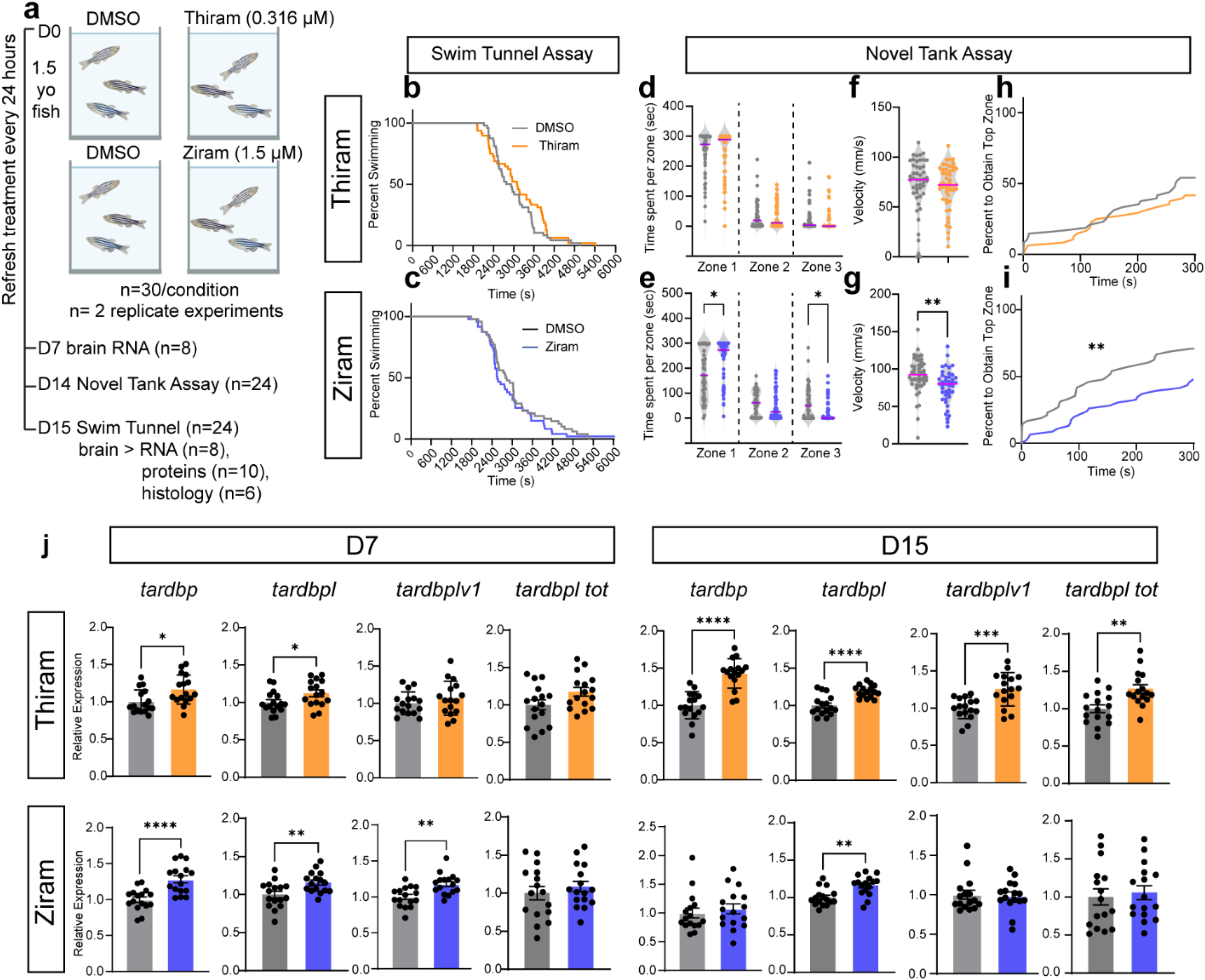
Exposure to thiram and ziram in vivo increases TDP-43 transcript levels and causes behavioral defects. **a** Experimental plan. **b-c** Swim tunnel assay results for thiram and ziram treatments expressed as percentage of fish swimming (y-axis) throughout the duration of the assay (x-axis, seconds). Black line = DMSO, colored line = treatment. **d-i** Novel tank assay results expressed as time spent per zone (sec) (**d-e**), total velocity (mm/sec) (**f-g**) and latency to enter zone 3 (top zone) in thiram (**d-f-h**) or ziram (**e-g-i**) exposures. Violin plots showing median (purple bar) and frequency of single fish measurements (dots) analyzed by Unpaired t-test. Survival plots data analyzed by Mantel-Cox test. n=48 fish per condition pooled from two replicate experiments. **J** RT-qPCR analysis of *tardbp*, *tardbplv1* and *tardbpl* variants and total *tardbpl* expression at day 7 and 15 of treatment with thiram (upper panel) and ziram (lower panel). Bars represent mean ± SEM of RNA levels relative to *ef1a* and normalized to DMSO controls; dots represent n=16 fish/condition. Unpaired t-test. *p < 0.05, **p < 0.01, ***p < 0.001, ****p < 0.0001. Only significant p values are shown. Further statistical information is available in the Statistical Data File. Extended data are available in the Extended Data File.

**Figure 6.**
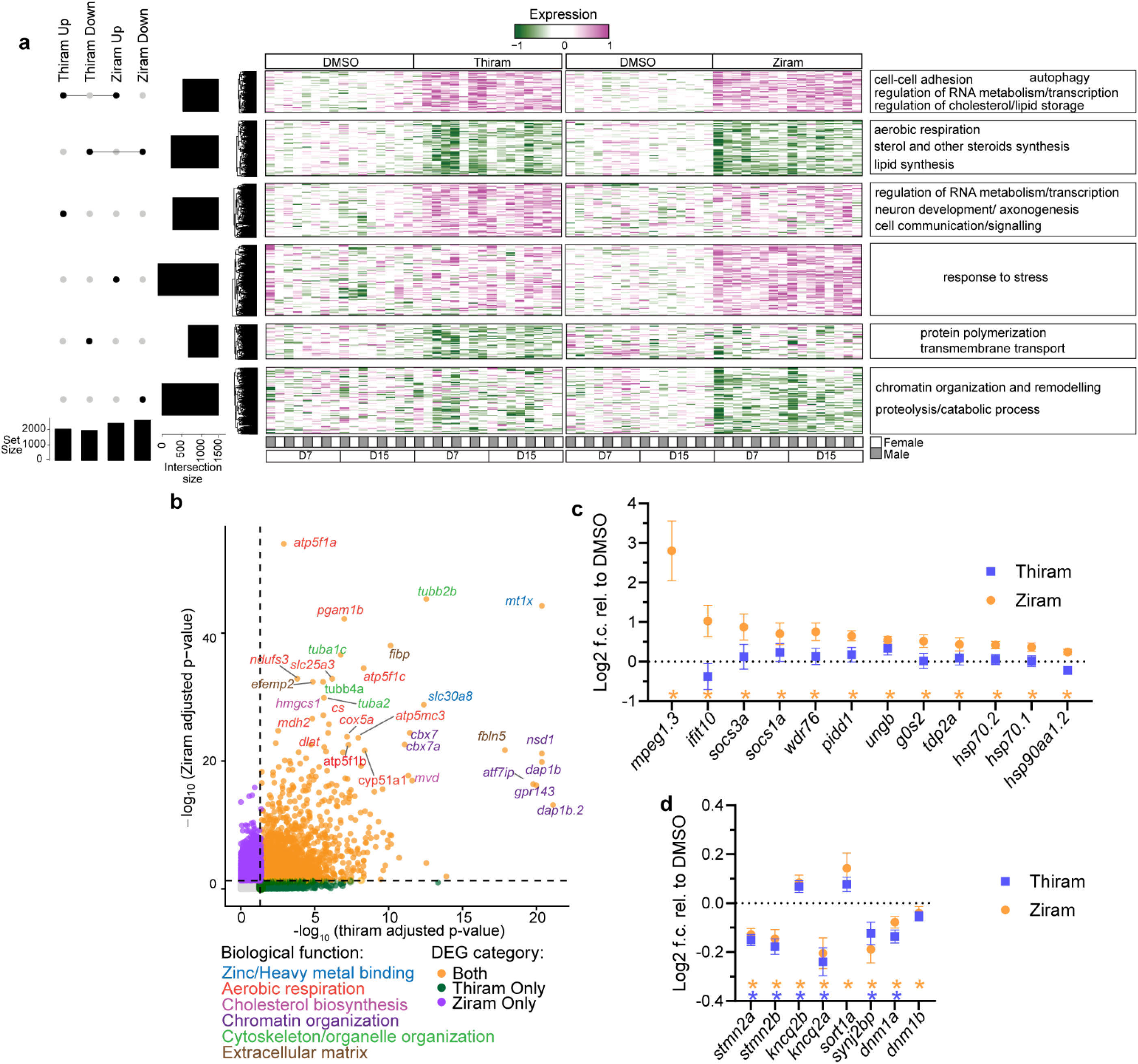
Exposure to thiram and ziram causes widespread transcriptional dysfunction. **a** Comparison of gene expression data from thiram and ziram treated fish brains identifies six clusters of differentially expressed genes, as represented by an UpSet plot (left). Heatmap shows hierarchical clustering of normalized gene expression values, centered around the mean of DMSO samples for each treatment (pink = higher than that of DMSO, green = lower than DMSO). Summary of Gene ontology enrichment analysis results are shown on the right of each cluster. n=8 fish (4 males +4 females) for each time point and treatment conditions. **b** Scatter plot showing comparison of the adjusted p-values of the differentially expressed genes identified in thiram and ziram treated fish transcriptional analyses. Orange dots are DEGs shared between the two treatments, green dots are thiram-only DEGs, purple dots are ziram-only DEGs. The top 30 shared DEGs are labelled and colored based on their main biological function. **c-d** Expression levels of selected genes extracted from transcriptional analysis of thiram and ziram treated fish. **c** shows the top 12 stress response genes upregulated in ziram treated fish brains. **d** shows fish orthologues of genes previously identified as TDP-43 targets. Symbols represent mean ± SD of log_2_ fold changes relative to DMSO condition; n=16 fish per condition. blue squares = thiram, orange circles = ziram. Asterisks indicate differential expression significance in the corresponding dataset: orange = ziram, purple = thiram. Extended data are available in the Extended Data File.

As a result of a gene duplication event, zebrafish have two *TARDBP* orthologs: *tardbp*, which produces a full-length protein (412 aa; 73% similarity to human TDP-43), and *tardbpl*, which predominantly generates a truncated protein (303 aa) lacking the C-terminus. Under conditions of *tardbp* loss of function, a compensatory alternative splicing event generates a near full-length *tardbpl* variant (398 aa) referred to as *tardbpl-v1* which is thought to compensate for the reduction in TDP-43 expression and/or function [53].

To assess whether DTC exposure alters any of the zebrafish TDP-43 orthologs, we isolated zebrafish brains after 15 days of DTC exposure, generated brain homogenates, and performed immunoblotting using well validated zebrafish-specific TDP-43 antibodies [54]. Neither the expression nor gel mobility of Tardbp or Tardbpl-v1 proteins were significantly altered following thiram or ziram treatment (Supplementary Fig. 10a), which was also confirmed by immunofluorescence analysis of the telencephalon in thiram treated fish (Supplementary Fig.10b, c).

To determine any effects at the transcriptional level, we performed qRT-PCR using primers for *tardbp,* the two transcript variants, *tardbpl* and *tardbpl-v1,* and all t*ardbpl* isoforms. The analysis revealed a thiram-dependent increase in *tardbp* and *tardbpl* at day 7, with all orthologues significantly upregulated by day 15 (Fig. 5j). In contrast, ziram treatment induced an initial increase in *tardbp, tardbpl* and *tardblv1* transcripts on day 7, and the elevated expression was only sustained for *tardbpl* by day 15 (Fig. 5j).

To characterize the global transcriptional response to DTCs, we performed bulk whole-brain RNA sequencing at days 7 and 15 of ziram or thiram exposure (Fig. 6). As sex and treatment duration had minimal impact on transcriptional profiles, we included these variables as covariates and focused on the chemical treatment as the main effect. We identified 3,974 and 5,014 differentially expressed genes (DEGs; adjusted p < 0.05) in thiram-and ziram-treated fish, respectively (Fig. 6 and Supplementary Files 1, 2).

The DEGs clustered into six distinct groups revealing substantial overlap in the response to thiram and ziram, with 2,081 DEGs showing a similar regulation (Fig. 6a, b and Supplementary Files 1, 2). Shared upregulated genes were enriched for cellular processes including cell–cell adhesion, RNA metabolism, lipid and cholesterol regulation, and autophagy (Fig. 6a, Supplementary File 2b). Notably, metallothionein-1 (*mt1x*) and *slc30a8*, both encoding zinc-binding proteins, were among the top DTC-induced genes, again supporting a role for zinc homeostasis as a mechanism of action (Fig. 6b), consistent with restored TDP-43 solubility that we observed in the presence of zinc chelators (Fig. 2k, l). Shared downregulated genes were enriched for pathways related to aerobic respiration and lipid/steroid biosynthesis (Fig. 6a, Supplementary File 2b). Additionally, multiple tubulin genes (*tubb4b, tubb5, tubb2b, tuba1a, tuba1c*) were consistently downregulated, suggesting chemical-induced cytoskeletal disruption (Fig. 6a, b and Supplementary File 2).

Ziram exposure elicited a more robust stress response compared to thiram, characterized by upregulation of immune-related genes (*mpeg1.3, ifit10, socs3a, socs1a*), DNA damage and apoptosis-associated genes (*wdr76, pidd1, ungb, g0s2, tdp2a*), and molecular chaperones (*hsp70.1, hsp70.2, hsp90aa1.2*) (Fig. 6a, c and Supplementary File 2).

To assess whether the transcriptional changes associated with DTC exposure reflected TDP-43 loss of function, we compared zebrafish DEGs with their mouse and human orthologues identified in prior established TDP-43 datasets (Supplementary File 3). These include data obtained from TDP-43 knockdown mouse striatum [12], TDP-43 negative neurons from human FTLD-TDP brains [55] and TDP-43^K145Q^ knock-in mice in which TDP-43 RNA binding function is disrupted [48].

Despite the limited cross-species conservation of TDP-43 target genes, we found several TDP-43-regulated zebrafish orthologous that were differentially expressed in response to thiram and ziram. Notably, *stmn2a* and *stmn2b*, orthologs of the canonical TDP-43 splicing target *STMN2*, were downregulated by both thiram and ziram, similar to what observed in FTLD-TDP brain (Fig. 6d). Zebrafish orthologs of *KCNQ2*, another well characterized TDP-43 splicing target, showed divergent opposing regulation with *kcnq2a* being downregulated and *kcnq2b* being upregulated. Zebrafish orthologs of the mouse TDP-43 targets *Synj2bp* and *Dnm1* (*dnm1a* and *dnm1b*) were both decreased in response to thiram and ziram (though *dnmn1b* reduction only met significance with ziram treatment) (Fig. 6d). A complete list of shared genes in the different datasets is provided in Supplementary File 3.

Together, these findings indicate that the DTC family members thiram and ziram recapitulate many of the key molecular features of TDP-43 dysfunction in vivo, while also revealing disease-related pathways that could reflect distinct environmental-driven signatures of TDP-43 proteinopathies.

## Discussion

In this study, by using unbiased high-content imaging from a curated library of over 1,000 EPA-prioritized compounds, we identified a set of environmental chemicals, including DTCs, that induce TDP-43 aggregation and perturb TDP-43 proteostasis. These findings bolster a stronger link between environmental exposures and neurodegeneration, driven by chemicals with a shared ability to disrupt TDP-43 homeostasis and downstream molecular pathways that are central to ALS, FTD, and other TDP-43 proteinopathies.

### DTCs affect cysteine reactivity and metal homeostasis

A central observation of this study is that DTC-induced TDP-43 aggregation is critically dependent on reactive cysteine residues, highlighting an intrinsic biochemical vulnerability of TDP-43. Some DTCs are known to be thiol-reactive compounds capable of modifying proteins with diverse cellular functions, including aldehyde dehydrogenase, brain glycogen phosphorylase, and E1 ubiquitin ligase [37, 43, 45, 46]. However, a direct connection to TDP-43 had not been established. Thiram and disulfiram led to TDP-43 intermolecular disulfide crosslinking, as determined by loss of free thiol groups, which led to the formation of high-molecular weight TDP-43 aggregates. Our cysteine blocking and mass spectrometry data indicate that these DTCs likely directly modify TDP-43 reactive cysteine residues. However, these compounds have pleiotropic cellular effects that can potentially perturb cellular redox balance through other indirect mechanisms. Indeed, the oxidative stress marker HO-1 was induced in thiram-treated neurons, indicating that a global oxidative stress response could further amplify TDP-43 dysfunction and create a vicious cycle that feeds back to further impair TDP-43.

Despite some DTCs directly modifying cysteines, ziram does not promote TDP-43 cysteine crosslinking directly (i.e., in a cell-free system) but nonetheless promotes cysteine-dependent TDP-43 aggregation in cells, supporting an indirect effect on TDP-43. Our data, in line with previous studies, indicated that ziram acts as a zinc ionophore, increasing intracellular zinc levels and promoting metal-induced oxidative stress [46], thus providing a plausible explanation for its more potent effects as compared to thiram. Indeed, transcriptional profiling in zebrafish revealed a pronounced induction of stress-response pathways following ziram exposure. Notably, however, both thiram and ziram altered genes involved in zinc transport, aerobic respiration and mitochondrial function, suggesting that disruption of cellular redox and metal homeostasis represents a shared cellular pathomechanism that impacts TDP-43. Prior studies showed that zinc directly interacts with TDP-43 to promote its aggregation [29, 56], in support of a metal homeostasis model that regulates TDP-43 stability and function. Na-Pyrithione, a zinc ionophore [57] used as a preservative in both personal care and industrial products, was also identified as a strong inducer of TDP-43 aggregation in our study. Together, these findings provide new insight into how environmental toxicants converge on TDP-43 through the disruption of cysteine redox balance and metal homeostasis, processes that could be exploited to suppress aberrant TDP-43 species that accumulate in response to oxidative stress.

### DTCs effect on TDP-43 function

Thiram and ziram led to splicing defects that are consistent with TDP-43 loss of function, manifested in the induction of CE inclusion in mouse and human neurons. These effects were reflected in vivo, where exposure led to transcriptional signatures that partially overlapped with prior TDP-43 loss-of-function datasets. Notably, the downregulation of *stmn2* orthologs and dysregulation of *kcnq2*, *synj2bp*, and *dnm1* family members in zebrafish brains supports the conservation of TDP-43-regulated pathways across evolutionarily distant species. We also noted clear differences between DTC family members, as ziram generated a more pleiotropic and widespread transcriptional response characterized by changes in stress, immune, and metabolic pathways, consistent with a global stress response that is both TDP-43 dependent and independent.

Among DTCs, our findings highlight both thiram and ziram as particularly potent modifiers of TDP-43 function. Ziram exposure has been associated with increased risk of Parkinson’s disease in epidemiological studies [44], supporting its potential for neurotoxicity. Thus, we speculate that a single chemical entity can give rise to distinct clinical manifestations depending on dose, duration, and genetic background, among other variables. Whether DTCs might also be linked to tau pathology is unclear, however the presence of multiple pathologies in the postmortem brains of Guam ALS–parkinsonism–dementia patients [25, 26, 58] suggest that environmental exposures are sufficient to drive co-occurring pathologies. One possibility is that the combination of DTC-driven oxidative stress, metal imbalance and aberrant cysteine adducts/modifications lead to general proteostasis failure that promotes the misfolding of aggregate-prone proteins and thereby contributes to the slow, gradual emergence of end-stage brain pathologies.

Despite the fact that widespread transcriptional alterations, including in TDP-43 orthologues genes, were observed, neither thiram nor ziram induced overt motor deficits within the exposure windows tested, suggesting that early molecular changes precede overt phenotypic manifestations, at least in the case of ALS-like motor deficits. These findings are consistent with the prolonged prodromal phase characteristic of TDP-43 proteinopathies [2] and support a model in which environmental exposures initiate subtle yet cumulative effects on neuronal function.

### Gene-environment interactions

Our findings support a model in which environmental exposures could interact with genetic vulnerability to modulate disease-relevant phenotypes. By inserting a genetic substitution into the *TARDBP* locus to simulate TDP-43 acetylation-dependent RNA-binding deficiency (K145Q), we were able to demonstrate additive sensitivity to thiram, consistent with a gene-environment interaction. We note that TDP-43^K145Q^ is not a disease-associated genetic mutation [59], and therefore additional studies are required to determine whether disease-linked ALS/FTD mutations (in *TARDBP* or other ALS/FTD genes) similarly enhance susceptibility to environmental exposures. This additive effect was less evident for ziram at higher doses, potentially reflecting a ceiling effect due to the heightened stress response observed with ziram. Nonetheless, these findings are consistent with a multi-hit hypothesis previously proposed for ALS [47] and suggest that environmental factors can synergize with genetic lesions to lower the threshold required for TDP-43 dysfunction. Indeed, DTC exposure paradigms in vitro and in vivo provide new opportunities for ALS/FTD model development to better understand how environmental chemicals may accelerate disease progression in genetically susceptible individuals.

### Limitations

Several limitations of this study should be mentioned. The primary chemical screen relied on a TDP-43 overexpression reporter model challenged with acute chemical exposures, which may bias detection toward strong, rapid aggregation phenotypes and limit the identification of chemicals that impact TDP-43 in a more subtle manner manifested after chronic, low-dose, exposure paradigms. Nonetheless, our experimental design enabled a robust and scalable platform to uncover compounds that directly perturb TDP-43. Second, by employing TDP-43 aggregation as a primary readout, we may have biased our discovery towards compounds that induce visible TDP-43 inclusions, rather than compounds that simply induce TDP-43 loss of function via alternative mechanisms, including nuclear TDP-43 clearing or TDP-43 dissociation from its bound RNA. Nonetheless, TDP-43 aggregation is a central hallmark of TDP-43 proteinopathies and provides a relevant anchor for candidate selection. Validation across primary mouse neurons, human iPSC-derived neurons, and zebrafish models underscores the robustness and translational relevance of the identified candidates. Finally, although the concentrations used in vitro may exceed typical human exposures, they reveal mechanistic pathways through which sustained or cumulative exposure could disrupt TDP-43 homeostasis. Notably, disulfiram, a DTC used in a clinical setting for the treatment of alcohol use dependence and more recently as an anticancer agent [35, 36, 60], shares key thiol-reactive and redox-modulating properties [39–42, 61, 62], raising the concern that prolonged exposure to disulfiram and similar agents may inadvertently drive TDP-43 dysfunction.

In summary, our study identifies DTCs and other chemicals as environmental triggers for TDP-43 dysfunction. These findings provide a framework to now include specific environmental exposures as dominant factors in disease etiology and support a model in which the environment drives susceptibility to ALS/FTD and other TDP-43 proteinopathies.

## Material and Methods

### High-content imaging screen of ToxCast phase I/II libraries and validation

ToxCast phase I/II libraries (ph1_v2 and ph2) were received in 96-well plates at a 20 mM concentration and redistributed to a total of 5×384 well-plates using a TECAN Freedom EVO liquid handler. 6-12 wells of DMSO (negative control) and 6-12 wells of arsenite (positive control) were spiked in each plate. iGFP-TDP43-dNLS HEK-293A cells [34] were plated at 4.5×10^5 cells/well in a total of 45 384-well clear bottom black wall plates (Greiner 07-000-046) in media containing DMEM (Corning 10-013-CV), 1× penicillin/streptomycin (Gibco 15140122), 1x L-glutamine (Gibco 25030081), 10% tetracycline-free FBS (Avantor 76308-946) supplemented with 0.5 µg/ml of doxycycline (Sigma-Aldrich D5207). 24-hours later, cells were treated with 3 different concentrations of the ToxCast chemicals (4-20-100 µM) in 3 replicates (on independent plates) for 6 hours. Plates were then fixed in 4% PFA containing 1% Triton X-100 and imaged at in the GFP channel with a GE IN Cell Analyzer (20X, 5 fields/well). A TECAN Freedom EVO liquid handler was used on every liquid handling step. Validation of the independently acquired chemical hits in iGFP-TDP43-dNLS and iGFP-TDP43-WT HEK-293A cells [34] was performed using a similar protocol by using an OT-2 Liquid Handler (Opentrons Labworks Inc). All ToxCast chemicals independently acquired (Supplementary File 1b, c) were diluted in DMSO at 100 mM concentration. Arsenite was diluted in water at 100 mM. All chemicals were aliquoted in single use vials and stored at-80°C for a maximum of 3 months.

### Image analysis pipeline for high-content screen and validation

Image analysis in iGFP-TDP43-dNLS and iGFP-TDP43-WT HEK-293A was performed using a customized pipeline in CellProfiler ver 4.2.7 [63, 64]. GFP positive nuclei were identified by using the IdentifyPrimaryObjects module on the GFP images after image intensity was rescaled and the aggregate signal was subtracted by using the RescaleIntensity and EnhanceOrSuppressFeatures modules. GFP positive nuclei were expanded of 40 pixels (Expand_nuclei_40) to identify the cytoplasm. GFP-TDP-43 aggregates were then identified using the IdentifyPrimaryObjects module after using the enhance Speckles feature of the EnhanceOrSuppressFeatures module on the GFP images. A similar approach was used to identify phospho-TDP-43 aggregates in the validation experiment. Aggregates were then assigned to whole cells (Expand_nuclei_40). For each field the following parameters were calculated: (1) total number of GFP positive cells, percentages of whole cells with GFP-TDP-43 or phospho-TDP-43 aggregates, total number of GFP-TDP-43 or phospho-TDP-43 aggregates and average number of GFP-TDP-43 or phospho-TDP-43 aggregates per cell.

### High-content screen statistical analysis

For each of the 45 plates, the median number of cells in all the vehicle fields was first calculated and used to calculate the percentage of cells per field relative to vehicle for every field. All fields with less than 30% of cells per field relative to vehicle were then excluded from further analysis.

For each of the above-mentioned parameters, the average field values were calculated to obtain a single value per well. Data from triplicate plates were pooled and mean values of all vehicle wells were calculated. No significant plate to plate variability was observed by comparing the mean values of vehicle wells for each parameter. Fold changes of the mean well triplicate values relative to the mean well triplicate values of vehicle were then calculated. p-values were then obtained by performing a two-tailed t-test followed by a Benjamini-Hochberg’s false discovery rate (FDR) adjustment using R [65, 66]. Data from each dose were then pooled and the average field value of each parameter was calculated.

### Mouse husbandry

Mice were housed in ventilated microbarrier cages on racks providing high efficiency particulate air (HEPA)-filtered air supply to each cage. Animals were kept on a 12-hr light–dark cycle with access to food and water ad libitum. All animal husbandry, experiments, and procedures were performed in strict compliance with animal protocols approved by the Institutional Animal Care and Use Committee (IACUC) of the University of North Carolina at Chapel Hill (Protocol #24.190).

### Mouse primary neuron cultures

Murine primary cortical neurons were extracted and cultured as previously described [59]. Briefly, wild-type C57Bl/6 mice (Charles River) or TDP-43 WT/K145Q breeding pairs. Timed pregnant females at embryonic days 15–16 were lethally anesthetized with isoflurane, embryos were extracted and placed in cold 4-(2-hydroxyethyl)-1-piperazineethanesulfonic acid (HEPES)-buffered Bank’s balanced salt solution (HBSS) for brain isolation. For TDP-43 WT/K145Q dissections, process was paused for approximately 2 hr to permit genotyping of the fetuses, during which the brains were stored at 4°C protected from light in a Hibernate-E (BrainBits NC9063748) solution supplemented with B27 (Gibco 17504044) or NeuroCult SM1 (STEMCELL 05711) and GlutaMAX (Thermo Fisher 35050061). After genotyping, if applicable, the cerebral cortices from each brain were isolated and digested for 30 min at 37°C in a filter-sterilized HBSS solution containing 20 U/ml papain (Worthington Biochemical LS003126), 1 mM EDTA, 0.2 mg/ml l-cysteine, and 5 U/ml DNAse (Promega M6101). The enzyme solution was removed, and the digested tissue was washed twice with sterile HBSS. Warm plating media [BrainPhys media (Stemcell 05790), 5% fetal bovine serum, 1× penicillin/streptomycin (Gibco 15140122), 1× B27, 1× GlutaMAX] containing 5 U/ml DNAse, was added and the tissue was dissociated mechanically using a P1000 pipette. The resulting cell suspension was spun down for 5 min at 1.5 rcf to pellet the cells, resuspended in plating media, and filtered through a 40-mm cell strainer. Cells were counted and plated onto poly-D-lysine (PDL)-coated 12-well tissue culture plates (Corning 356470) at 300 K cells/well (for RNA and protein extraction) or 96-well glass bottom black wall plates (Cellvis P96-1.5H-N) at 30 K/well or 384-well clear bottom black wall plates (Greiner 07-000-046) at 15 K/well (for immunofluorescence and microscopy). 16–24 hr after plating, all plating media was removed and replaced with neuronal cell media (BrainPhys, 1× GlutaMAX, 1× B27 or SM1, Glucose 15mM, 1× penicillin/streptomycin). Cultures were incubated at 37°C, 5% CO_2_ and 95% humidity with half-media exchanges every 3 days for the duration of all experiments.

### hSyn-GFP-TDP43-dNLS construct generation and lentivirus preparation

Lentiviral vectors were constructed in-house using the pUltra lentiviral vector backbone for cloning via restriction enzyme digestion and ligation. Briefly, the pUltra construct was acquired from Addgene (plasmid # 24129) and the EGFP-P2A was substituted with an EGFP construct; The TDP-43-dNLS sequence [67] was then inserted downstream EGFP into the pUltra vector backbone. The human synapsin (hSyn) promoter was PCR amplified using primers listed in Supplementary Table 2. The resulting PCR product was used to replace the human ubiquitin C promoter in pUltra-GFP-TDP43-dNLS via PacI and AgeI digestion and ligation, generating pLV-hSyn-GFP-TDP43-dNLS. The resulting ligation products were transformed into NEB Stable competent cells using standard protocols. Lentiviral production and purification was performed following standard procedures as previously described [59].

### hSyn-GFP-TDP43-dNLS infection of mouse cortical neurons, chemical treatment and quantification of TDP-43 aggregation

E16 mouse cortical neurons were plated in 3 384-well plates and infected with the hSyn-TDP43-dNLS-GFP lentivirus on day in vitro (DIV) 8. On DIV 13, when over 90% of the TDP43-GFP signal was localized in the cytoplasm, neurons were treated with the 14 chemicals hits identified in the screen and listed Supplementary File 1c. Arsenite was used as positive control and DMSO as negative control. Each chemical was used at 5 different concentrations (0.16, 0.8, 4, 20, 100 µM) in duplicate wells for 3.5 hours. Propidium iodide (PI, Sigma Aldrich P4864) was added (1 μg/mL) 1.5 hours after the chemical treatments. Cells were then fixed in 4% PFA, washed four times with 1× PBS, with the second wash containing 1 µg/ml 4′,6-diamidine-2′-phenylindole dihydrochloride (DAPI). Plates were preserved in an 85% glycerol in 1× PBS solution at 4°C. Cell were imaged in the GFP/PI/DAPI channels on a GE IN Cell Analyzer 2000 at 20x, 4 fields/well for a total of 8 fields per condition. Image analysis was performed on CellProfiler [63, 64]. Live neurons were identified based on nuclear size and level of PI signal in nucleus, by filtering out bright PI nuclei (dead cells) and smaller nuclei (non-neuronal cells). The PI signal in the cytoplasm of live neurons was used to define the neuronal cytoplasm. TDP43-GFP positive neurons were identified by setting a threshold on the TDP43-GFP signal in the cytoplasm of live neurons. TDP43-GFP Granularity in TDP43-GFP+ live neurons was measured using the MeasureGranularity function. Three iterations of the MeasureGranularity module were selected as best representative of the amount of granularity in the positive control (arsenite) and used to measure TDP43-GFP granularity in the other conditions. Fold change of viability and TDP43-GFP granularity were calculated by dividing the number of live neurons and granularity in every condition to the geometric mean of the number of live neurons in the negative control conditions in the same plate. Statistical significance was calculated using a Krustal-Wallis non-parametric test with false discovery rate correction (n=8 fiels/condition; DMSO 0.00016% used as control).

### Solubility fractionation and immunoblotting

All steps of protein fractionation were performed on ice unless otherwise indicated. For each well of a 12-well plate, cells were suspended in 150 µl RIPA buffer (50 mM Tris pH 8.0, 150 mM NaCl, 1% NP-40, 5 mM EDTA, 0.5% sodium deoxycholate, 0.1% sodium dodecyl sulfate [SDS]) containing a mix of protease, phosphatase, and deacetylase inhibitors [1 µg/ml Peptstatin A (Sigma P4265), 1 µg/ml Leupeptin (Sigma L2023), 1 µg/ml Nα-Tosyl-l-lysine chloromethyl ketone hydrochloride (TPCK) (Sigma T7254), 1 µg/ml Trypsin inhibitor (Sigma T9003), 1 µg/mL N-p-Tosyl-l-phenylalanine chloromethyl ketone (TLCK) (Sigma T4376), 0.67 µg/ml trichostatin A, 10 mM nicotidamide, 1 mM phenyline thanosulfyl fluoride, 1 mM phenylmethylsulfonyl fluoride]. The solution was homogenized by sonication and centrifuged at 4°C for 45 min at 18,000 × rcf. The supernatant was removed and saved as the RIPA-soluble (soluble) protein fraction. The pellet was resuspended in 500 µl RIPA buffer with inhibitor mixture, sonicated, and centrifuged as described above, and the supernatant discarded. The resulting pellet of RIPA-insoluble material was resuspended in 75 µl of Urea buffer (7 M urea, 2 M thiourea, 4% CHAPS, 30 mM Tris, pH 8.5) with the inhibitor mixture (as above), sonicated to homogenize, and centrifuged at room temperature (RT) for 45 min at 21,000 × rcf. The resulting supernatant of RIPA-insoluble, urea-soluble protein fraction was saved as the ‘insoluble’ protein fraction.

Equal volumes of samples were run onto 4-20% Tris-Glycine SDS-polyacrylamide gel electrophoresis gels (Bio-Rad 5671095) in the presence or absence of dithiothreitol (DTT, Sigma D0632) and then transferred onto nitrocellulose membranes. Membranes were washed three times in Tris-buffered saline with 0.1% Tween-20 (TBST), once in 1x Tris-buffered saline (TBS), and blocked in 4% nonfat milk in 1× TBS for 1 hour at RT. Membranes were incubated with primary antibodies (Supplementary Table 1) diluted in 4% milk overnight at 4°C. The primary antibody solution was removed and membranes washed three times in TBST, once in TBS, and incubated with cross adsorbed horseradish peroxidase (HRP)-conjugated goat secondary antibodies (Supplementary Table 1) diluted in 4% milk at 1:5,000 for soluble or insoluble protein immunoblots, respectively, for 3 hr at RT. HRP signal were detected using either Pierce ECL Western Blotting Substrate (Thermo Scientific PI32106) for more abundant proteins or Amersham ECL Select Western blot detection reagent (Cytiva RPN2235) for lower abundance proteins. Blots were then visualized by chemiluminescent imaging using an ImageQuant 800 Fluor (Cytiva) machine. If multiple antibodies were used on the same membrane, signal was stripped using 1X ReBlot Plus Strong Antibody Stripping Solution (Millipore Sigma 2504). All reagents were used following manufacturer instructions. Densitometry analysis to quantify western blot images was performed in LI-COR Image Studio Lite (Lincoln, NE, USA). Protein amounts were normalized to the loading controls specified in each experiment.

### rTDP-43 chemical incubations and quantification of TDP-43 species

rTDP-43 (R&D Systems AP-190) was incubated with DMSO or 200 µM of the different chemicals (thiram, ziram, disulfiram, SDTC, Na-pyrithione and BIT) at RT for 1 hour (0.5 µg rTDP-43 /reaction), followed by labeling with 80 µM of 5-iodoacetamidofluorescein (5-IAF, Sigma Aldrich I9271) for 10 minutes at 37°C. Reactions were then aliquoted in two tubes and separated by Sodium Dodecyl Sulfate Polyacrylamide Gel Electrophoresis (SDS-PAGE) in the presence or absence of DTT and transferred onto nitrocellulose membranes. 5-IAF fluorescence was directly detected from the +DTT membrane using an ImageQuant 800 Fluor imager (Cytiva) while rTDP-43 signal was measured by chemiluminescence after incubation with the TDP-43 3H8 antibody (Supplementary Table 1). Image Studio Software (LI-COR Biotech, LLC) was used to quantify the fluorescence and chemiluminescence signal in western blot membranes. Total rTDP-43 in - DTT blots were obtained by quantifying the signal in a rectangle area spanning all rTDP-43 species (top of the well to 37 kDa). Signals from monomeric (43 kDa), high molecular weight (>250 kDa), trimeric (∼130 kDa) and tetrameric (∼170 kDa) rTDP-43 were quantified and divided by the total rTDP-43 signal to obtain the percentages of each species. The percentage of rTDP-43 in the smear (oligomeric) was obtained by subtracting the percentages of the different species from the total. 5-IAF signal in +DTT blots was normalized to total rTDP-43 in +DTT membranes.

### Identification of cysteine crosslinks via LC-MS/MS analysis

#### Sample preparation

rTDP-43 (R&D Systems AP-190) was purified using Zeba™ Spin Desalting Columns 7 MWCO (Thermo Fisher Scientific 89890) as per manufacturer instructions, and resuspended in 350 µl of 100 mM ammonium bicarbonate (ABC) buffer (Millipore Sigma S2454). rTDP-43 protein concentration was then quantified using NanoDrop 2000 spectrophotometer and aliquoted in single reactions tubes containing 25 µg of protein in 45 ul of ABC buffer. Purified rTDP-43 aliquots were incubated with DMSO, thiram (20 µM) or disulfiram (20 µM) for 1 hour, followed by incubation with 34 mM of chloroacetamide (Thermo Fisher Scientific A39270) for 10 minutes. Samples were separated by SDS-PAGE and stained with Comassie (Bio-Rad 1610786) overnight. Gel regions corresponding to monomeric (∼43 kDa) or HMW (>250 KDa) rTDP-43 were isolated for each sample and processed for LC-MS/MS analysis. To preserve putative cysteine cross-links created by TDP-43 treatment, no further reduction and alkylation was performed before digestion. Gel slices were first washed with 100 µL 1:1 solution of 100 mM ABC and acetonitrile (ACN, Thermo Fisher Scientific 615140025) then dehydrated with 500 µL ACN. Residual ACN was evaporated via centrifugal vacuum evaporation for about 5 min. A 20 µg vial of trypsin protease (Thermo Fisher Scientific 90059) was reconstituted into 1 mL of 9:1::10 mM ABC:ACN. A 100 µL aliquot was added for a total of 2 µg trypsin per gel band. Samples were incubated overnight in a 37°C oven. Digestion was quenched by adding 1:2::5% formic acid (FA, Theremo Fisher Sientific 28905) in water:ACN with a 15 minute ambient incubation and the supernatant transferred to clean microcentrifuge tubes. Gels were alternatingly dehydrated with 50 µL ACN and rehydrated with 50 µL 100 mM ABC 3 times. All supernatants were combined before evaporating completely by centrifugal vacuum evaporation. Peptides were reconstituted in 50 µL 98:2::water with 0.1% FA:ACN and transferred to LC autosampler vials containing a low-volume insert.

#### LC-MS/MS analysis

A 2 µL injection was analyzed by reversed phase nano-liquid chromatography-mass spectrometry (nanoLC-MS/MS) using a Vanquish Neo nanoLC system (Thermo Scientific, San Jose, CA, USA) interfaced with an Orbitrap Eclipse (Thermo Scientific) mass spectrometer. The ‘trap and elute’ configuration consisted of a 0.075 mm × 20 mm Accclaim PepMap^TM^ 100 C_18_ trap column with particle size of 3 µm (Thermo Scientific) in line with a 0.075 mm × 250 mm PepMap^TM^ C_18_ nanoLC analytical column with particle size of 2 µm (Thermo Scientific). Peptides were eluted using a solvent gradient of Mobile Phase A (MPA) containing water with 2% acetonitrile and 0.1% formic acid, as well as Mobile Phase B (MPB) containing acetonitrile with 20% water and 0.1% formic acid (MPB). MPB was held at 5% for 2 min, increased to 25% over 43 min, increased to 50% over 15 min, increased to 95% in 5 min, and was held at 95% for 8 min for a total run time of 75 min. Mass spectrometer parameters were set as follows: 1.8 kV positive ion mode spray voltage, ion transfer tube temperature of 275 ℃, master scan cycle time of 2 s, m/z scan range of 375 to 1,500 at 120,000 _FWHM_ resolving power (at m/z 200), 250% normalized AGC Target, 50 ms maximum MS^1^ injection time, RF lens of 40%, 15,000 _FWHM_ resolving power (at m/z 200) for data-dependent MS^2^ scans, 1.6 m/z isolation window, 30% normalized HCD collision energy, 250% normalized AGC Target, automated maximum injection time and dynamic exclusion applied for 90 s periods.

#### Data Interrogation

Raw nanoLC-MS/MS files were processed with MassMatrix Xtreme 3.0.10.25 software (MassMatrix, Columbus, OH) using the human TDP-43 protein sequence (414 amino acids, Taxon 9606)[68–71]. Results were tested against a reversed decoy sequence for TDP-43. Trypsin was designated as the cleavage enzyme hydrolyzing at lysine and arginine with proline as an inhibitor. Variable modifications were acrylamide adduct of cysteine, deamidation of asparagine and glutamine, carbamidomethylation of cysteine, and oxidation of methionine. Mass tolerance for MS^1^ and MS^2^ were 5 ppm and 0.02 Da, respectively. Disulfide cross links were searched using exploratory mode with disulfides non-cleavable by the enzyme and a maximum of 2 possible cross links per peptide. Allowable length for peptides was 6 to 60 amino acids. Data output required a minimum pp score of 5.0 and pp-tag score of 1.3.

### hiPSC-derived cortical neurons

Generation of TDP-43 K145Q hiPSCs, hiPSCs maintenance and cortical neurons differentiation have been previously described [59]. Mature cortical neurons were obtained using a modified dual SMAD inhibition protocol [72, 73]. Briefly, undifferentiated iPSCs were collected with Accutase (Thermo Fisher A1110501), counted and cultured in StemScale PSC medium (Thermo Fisher A4965001) supplemented with 10 µM Y27632 (PeproTech 1293823) and cultured in suspension on an orbital shaker at 37°C and 5% CO_2_. After 48 hr, cells were dissociated with Accutase and differentiated into neuronal precursor cells (NPCs) as neutrosphere in StemScale PSC medium supplemented with 10 µM Y27632 (PeproTech 1293823) and 1.5 µM CHIR99021 (PeproTech 2520691), 10 µM SB431542 (PeproTech 3014193) and 50 nM LDN-193189 (Sigma SML0755). CHIR99021 was removed after 24 hr and cells were cultured for 10 days with daily medium changes and neurospheres dissociated at 1:3 ratio twice a week. Then NPCs were expanded for 7 days in the presence of 20 ng/ml FGF (Peprotech 100-18B) in Neuronal Expansion Medium [1:1 Advanced Dulbecco’s Modified Eagle Medium (DMEM)/F12 (12634028) and Neural Induction Supplement (A1647801)]. NPCs were plated for maturation on PDL/Laminin-coated plates or coverslips in cortical neuron maturation medium [1:1 Advanced DMEM/F12 and Neurobasal; 1× GlutaMAX, 100 mM B-mercaptoethanol, 1× B27, 0.5× N2, 1× non-essential amino acids (NEAA), and 2.5 mg/ml insulin (Sigma-Aldrich I9278)] supplemented with 10 ng/ml BDNF, 10 ng/ml glial cell-derived neurotrophic factor (GDNF), and 10 µM N-[N-(3,5-difluorophenacetyl)-L-alanyl]-S-phenylglycine t-butyl ester (DAPT). Seventy-two hours after plating, cells were treated with 1 µg/ml Mitomycin C for (Sigma M5353) for 1 hr.

### Primary mouse neuron and hiPSC-derived cortical neuron chemical treatments

On DIV14 (mouse primary neurons) or mature hiPSC-derived cortical neurons (∼DIV50), half of the media was removed, and neurons were treated with one volume of fresh media containing 2X concentrated chemicals or DSMO for 3 hours. Cell viability in primary neurons experiments was measured by adding CellTiter-Blue Cell Viability Assay (Promega G8080) 30 minutes after starting the chemical treatments and incubated for 2.5 hours. Fluorescence was recorded using 560_Ex_ /590_Em_ on a plate reader. For pretreatment experiments, triethylenetetramineQ hydrochloride (TET, Sigma Aldrich 161969), N,N,N′,N′-tetrakis(2-pyridylmethyl)ethylenediamine (TPEN, Sigma Aldrich P4413), N-acetyl-L-cysteine (NAC, Sigma Aldrich A9165) or zinc (Sigma Aldrich Z0152) were added directly to the neurons for 40 minutes, followed by toxicant treatments as described above. For mouse experiments, biological replicates were defined as neuron cultures extracted from 3 independent embryos of each genotype (WT and TDP-43^K145Q^). For hiPSC-derived neurons experiments, biological replicates were defined as 3 independent neuron differentiations from the same hiPSC line, one line for each genotype (WT and TDP-^43K145Q^)

### Zebrafish Husbandry

Wild type AB fish were maintained at 28° C at the University of North Carolina Zebrafish Aquaculture Core Facility with a 14/10-hour light/dark photo period. All fish husbandry, anesthesia, euthanasia and experiments were approved by the IACUC of the University of North Carolina at Chapel Hill (Protocol #26-020.0)

### Zebrafish Chemical Exposures

Wild type AB fish were exposed for up to fifteen days to one of two chemicals, thiram or ziram. Fourteen-day dose response studies were previously conducted to determine the lowest observable adverse effect level (NOAEL) for each chemical (0.316 µM thiram, 1.5 µM ziram) to be used in main exposure trials. For main exposure trials, fish were selected from age-matched clutches and randomly assigned to one of two conditions, DMSO control or chemical. Fish for two separate thiram experiments were 14 and 15 months old, respectively. Fish for two separate ziram experiments were 17 and 18 months old, respectively. Fish were maintained in two liters of static fish system water spiked with DMSO or chemical. Fifty percent water changes were conducted daily, and water quality was monitored weekly by zebrafish core staff (total ammonia-N, nitrite, nitrate, pH, and conductivity). Exposed fish were fed once daily following water change with Skretting Gemmo Micro 300 (Nutreco, Norway). Fish were monitored and videoed daily.

### Zebrafish Novel Tank Assay

A novel tank assay based on a previously described protocol [74], was conducted to test anxiety-like behavior using the Zantiks LT system (Cambridge, UK) at fourteen days of exposure. Briefly, individual fish were taken from group tanks (two liters of water in a standard 3.5-liter tank) and introduced into a novel tank (13 cm x 36 cm x 14 cm). Movement was tracked and recorded for five minutes following entry into the arena. The arena was subdivided into three approximately equal vertical zones (zone 1, bottom; zone 2, middle; zone 3, top). Time spent and distance traveled per zone were recorded in one second bins.

### Zebrafish Swim Tunnel Assay

Swim endurance was tested against an incrementally increasing flow rate in a Loligo System swim tunnel (5–185 L) with an SEW-Eurodrive 60 Hz motor and SEW-Eurodrive 0.5 HP/0.37 kW Movitrac LTE controller (Loligo Systems, Viborg, Denmark). Twelve to fifteen fish were introduced into the swim chamber with a starting temperature of 26.5°C (±1°) and allowed to acclimate for twenty minutes at a flow rate of 7 cm/s (5 Hz motor output). Following acclimation period, motor output was increased to 6 Hz (10 cm/s) and increased by 1 Hz every two minutes, up to 60 Hz (151 cm/s). Fish were allowed to swim against the increasing current until they fatigued and were no longer able to maintain swimming position. Fatigued fish were immediately removed from the swim chamber, and individual swim time was recorded.

### Zebrafish Brain Tissue Dissection

Following completion of exposure experiments, fish were euthanized in ice-cold water for ten minutes per IACUC protocol. Brains were immediately removed and placed in 1.7 mL Eppendorf tubes as follows: for RNA isolation-in 300 µL TRIzol (Invitrogen 15596018), flash frozen in liquid nitrogen, and stored at-80°C; for protein extraction-in an empty tube, flash frozen in liquid nitrogen, and stored at-80°C; and for histology-in 300 µL 4% PFA, stored at 4°C for twenty-four hours, then washed and cryoprotected in 30% sucrose (in PBS) at 4°C.

### RNA Isolation

Zebrafish brains previously frozen in 300 µL TRIzol were thawed and homogenized with a pellet pestle, followed by addition of 350 µL TRIzol and 200 µL chloroform. For experiments involving mouse primary cortical neurons or human iPSC-derived cortical neurons, cells were washed with 1× PBS, harvested from 12-well plates using 500 µL TRIzol/well and stored at-80°C. Samples were thawed and let at RT for 5 minutes, followed by addition of 50 µL of 1-bromo-3-chloropropane (Acros Organics 106862500). Samples were mixed and incubated for 5 minutes at RT. Samples were centrifuged 12,000 rcf at 4°C for 15 minutes. The aqueous phase containing RNA was removed and mixed with one volume of 70% ethanol. Mixture was transferred to a Qiagen RNeasy spin column, and RNA was isolated per Qiagen RNeasy Plus (Qiagen, Inc. 74134) protocol using Qiagen RNase-free DNase1 (Qiagen, Inc 79254) for genomic DNA digestion. RNA was eluted in 30 µL RNase-free water and RNA concentration was quantified by NanoDrop spectrophotometer (Fisher Scientific, MA, USA).

### Quantitative Reverse Transcription-Polymerase Chain Reaction

cDNA was generated from 250 to 1000 ng of RNA using High-capacity RNA-to-cDNA kit (Applied Biosystems 4387406) per the manufacturer’s instructions. Quantitative PCR was performed using PowerUp SYBR Green master mix (Applied Biosystems A25776) on a QuantStudio 6 PRO Real Time Polymerase Chain Reaction system and analyzed using with Thermo Fisher Design & Analysis Software (version 2.7.0). The PCR phase consisted of 40 cycles of 15 s at 95°C and 1 min at 60°C. Primers for all the transcripts tested are listed Supplementary Table 2. cDNA amounts ranging from 0.5 to 20 ng were used per reaction, depending on the transcript abundance. *Gapdh*, *RPLP0* and *ef1a* were used as the reference genes in mouse neurons, hiPSC-derived neurons and zebrafish brains respectively. Primers were synthesized by Integrated DNA Technologies (IDT).

### Zebrafish Histology and analysis

Frozen zebrafish brains were embedded in optimal cutting temperature compound (23-730-571 Fisher HealthCare) and the telencephalon was sectioned coronally at 10 um and mounted on slides serially and frozen at-80°C. Slides were defrosted at RT for 1 hour, dehydrated in 3 sequential 2 minutes washes of deceasing percentages of ethanol solutions (100-95-70%), hydrated in 1XPBS for 10 minutes, permeabilized and blocked in 10% normal donkey and goat serum in TBS/TX (0.05M Tris, 2.7% NaCl, 0.3% Triton-X100, pH 7.6) for 1 h at RT. Tissue was then incubate in primary antibody in above blocking solution overnight at 4°C followed by 3 washes in TBS/TX and 2 hour incubation in secondary antibody mix in blocking solution. Slides were rinsed in 1XPBS and mounted using Fluoromount-G with DAPI (Invitrogen 00-4959-52). 3D images for Tardbpl and DAPI were collected at 63X on a Zeiss LSM 800 confocal microscope. Quantification of Tardbpl cell intensity and foci counts/cell on 3D images were performed using the ImageJ plugin Foci Analyzer (BioImaging Facility at the Netherlands Cancer Institute, Amsterdam).

### Zebrafish RNA-seq library preparation and analysis

RNA isolation was performed and quantified as previously described. RNA was then diluted to 50 ng with RNase-free water up to a final volume of 25 µL and submitted to the UNC High-Throughput Sequencing Facility. RNA was further quantified with both Qubit (Thermo Fisher Scientific) and TapeStation Analysis Software 4.1 (Agilent Technologies). RIN values ranged from 9.3-9.6. A custom library prep was conducted with Zymo Ribo-Free Total RNA Library Prep and QAQC with pooling. Sequencing was performed as T7 Paired End read type with 2 x 100 read length. RNA sequencing data from zebrafish exposed to thiram and ziram were processed using a standardized analysis pipeline implemented in Linux and Bash, with downstream statistical analyses performed in R. Transcript-level quantifications were generated using Salmon [75], and transcript abundances were summarized to gene-level counts using the tximport package [76]. Gene annotation and transcript-to-gene mappings were obtained from the Danio rerio reference genome GRCz11 using Ensembl gene annotations. All downstream analyses were performed in R using Bioconductor packages.

Differential gene expression analyses were performed using DESeq2 [77]. Separate analyses were conducted for thiram and ziram exposures. For the full dataset, differential expression between treatment and DMSO controls was modeled using a design formula that accounted for both sex and exposure duration (Day 7 and Day 15), using the model ∼ sex + duration + treatment. To evaluate time-specific effects, additional stratified models were performed within each duration using the design ∼ sex + treatment. Wald tests were used to estimate log2 fold changes and statistical significance, and p-values were adjusted for multiple testing using the Benjamini–Hochberg false discovery rate procedure. Genes with an adjusted p-value (padj) ≤ 0.05 were considered significantly differentially expressed.

For visualization and downstream comparisons, normalized gene expression values were generated using the variance stabilizing transformation (VST) implemented in DESeq2. To isolate treatment-associated transcriptional changes, batch effects associated with sex and exposure duration were removed from the VST-normalized expression matrix using the removeBatchEffect function from the limma package [78]. For visualization purposes, expression values were centered relative to the mean expression of DMSO control samples for each gene.

To compare transcriptional responses between thiram and ziram exposures, differentially expressed genes from each analysis (padj ≤ 0.05) were merged and classified according to their direction and significance in each dataset. Genes were categorized based on whether they were significantly upregulated or downregulated in thiram only, ziram only, or in both treatments. Intersections between these DEG sets were visualized using an UpSet plot generated with ComplexHeatmap [79], which quantified the number of genes shared between compound-specific and shared transcriptional responses.

To visualize expression patterns across treatments, a combined heatmap was generated using genes that were significantly differentially expressed in either Thiram or Ziram analyses. Expression values were log-transformed, batch-corrected for sex and duration, and centered relative to control samples. To enable direct comparison of expression magnitude between genes, expression values were scaled by the maximum absolute value observed for each gene. Heatmaps were generated using ComplexHeatmap, with samples annotated by compound exposure, treatment condition, duration, and sex.

To evaluate concordance between the observed transcriptional responses and previously reported TDP-43-associated transcriptional signatures, DEG lists from the thiram and ziram analyses were compared with published datasets derived from TDP-43 loss of function datasets. Zebrafish orthologues of human and mouse genes in these datasets were extracted using Ensembl BioMart. Ensembl IDs in the different datasets were intersected using R.

Gene ontology enrichment analyses were performed using gProfiler2 [80]. Enrichment testing was conducted separately for gene sets derived from the differential expression analyses and from DEG overlap categories. GO biological process terms were considered significant using the default multiple testing correction implemented in g:Profiler. All analyses and visualizations were performed in R using packages including DESeq2, tximport, limma, ComplexHeatmap, EnhancedVolcano, biomaRt, and tidyverse.

## Statistical Analysis

Statistical analysis for the high-content screen was performed in R (www.R-project.org). All other data were analyzed in GraphPad Prism version 10 (GraphPad Software, San Diego, CA, USA, https://www.graphpad.com). Details regarding the statistical test performed, sample sizes, and what the datapoints and error bars represent can be found in the appropriate figure legends. All statistical tests were two sided. All image analysis was blinded. No statistical method was used to predetermine sample size. Statistical significance was determined as p < 0.05. Supplementary statistical information about data presented in Figures and Supplementary Figures can be found in the Extended Data and Statistical Data files.

## Data Availability

RNA-Seq data have been deposited in the NCBI’s Gene Expression Omnibus (GEO) database with accession number GSE341813.

## Supporting information

Statistical Data File

## Acknowledgments

This study was supported by the NIH R01AG087877, the National Institute of Environmental Health Sciences of the National Institutes of Health under Award Number P30ES010126. This work was performed in part by the Molecular Education, Technology and Research Innovation Center (METRIC) at NC State University, which is supported by the State of North Carolina.

We would like to thank Gary AB Armstrong for donating the zebradish tardbpl and tardbplv1 antibodies. High-content imaging and confocal imaging of zebrafish sections were performed at the Hooker Imaging Core at UNC.

## Author Contributions

G.F, T.J.C. and A.P. conceptualized the study and obtained funding, G.F and T.J.C. wrote the manuscript and all other authors edited and approved it. G.F., R.D.W, J.W, A.F.B, K.N.K, V.B., O.K.A, A.B. X.T., A.J.E., J.C.K and B.A.E performed experiments and/or analyzed data and contributed and/or developed methodology, G.F., R.D.W, T.A.B, M.J.N, A.H, L.B.C and T.I.W analyzed data and performed data visualization, A.P., T.J.C. provided supervision. We would like to thank Gary Armstrong

---

**Supplementary Figure 1.**
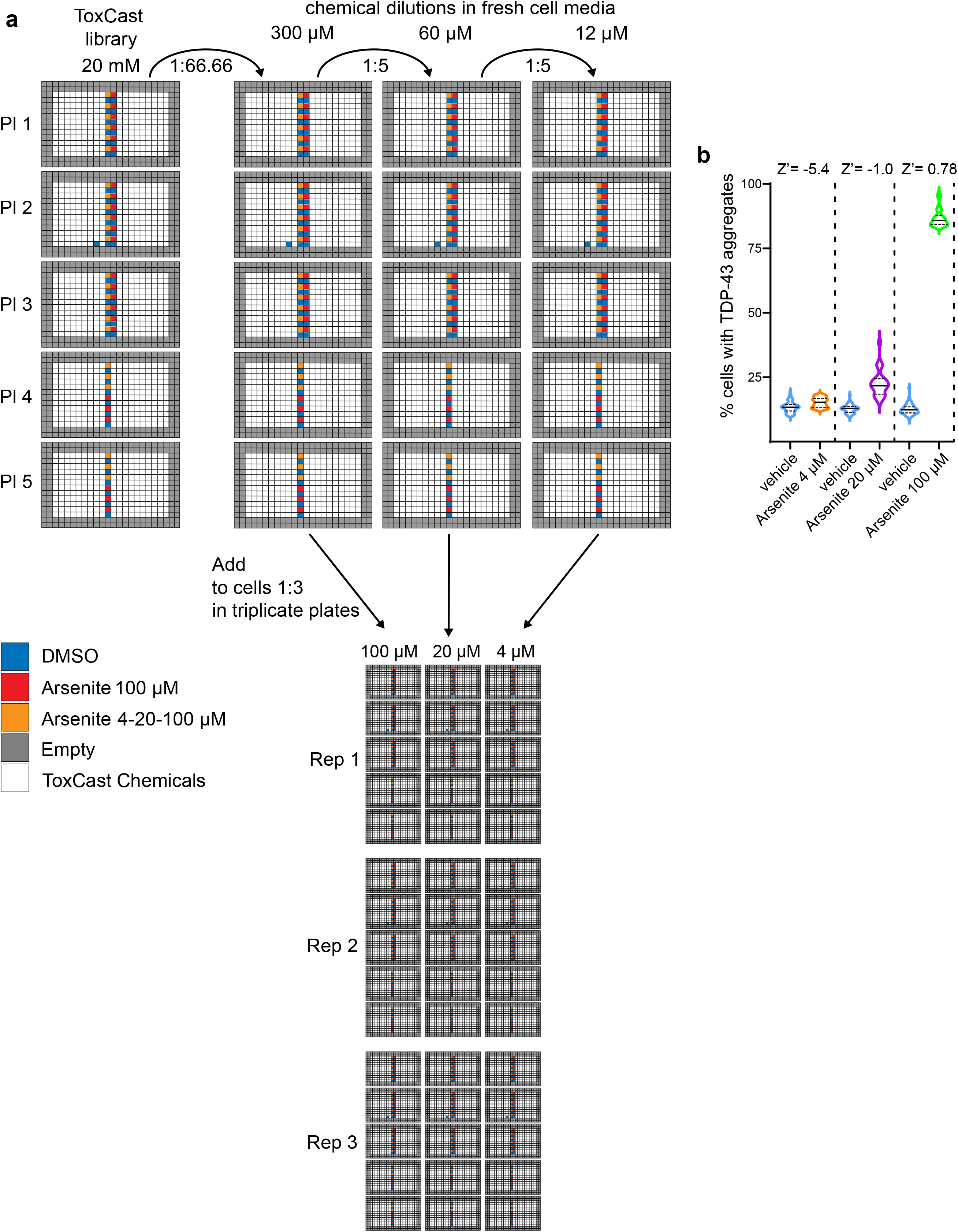
Screen plate setup. **a** ph1_v2 and ph2 ToxCast libraries were received in 96 well plates at a 20 mM concentration and redistributed to a total of 5 x 384-well plates using a TECAN Freedom EVO liquid handler. 6-12 wells of DMSO (negative control) and 6-12 wells of arsenite (positive control) were added on each plate. **b** Z-factor was calculated using average and standard deviation of all vehicle (n=55) and arsenite treated (n=24) wells in each set of screening plates (4,20 and 100 µM). Violin plots show averages of replicate wells of the % of cells with aggregates in vehicle and arsenite treated conditions. Dotted lines separate the data from the 3 different screen doses (4,20 and 100 µM). Further statistical information is available in the Statistical Data File. Extended data are available in the Extended Data File

**Supplementary Figure 2.**
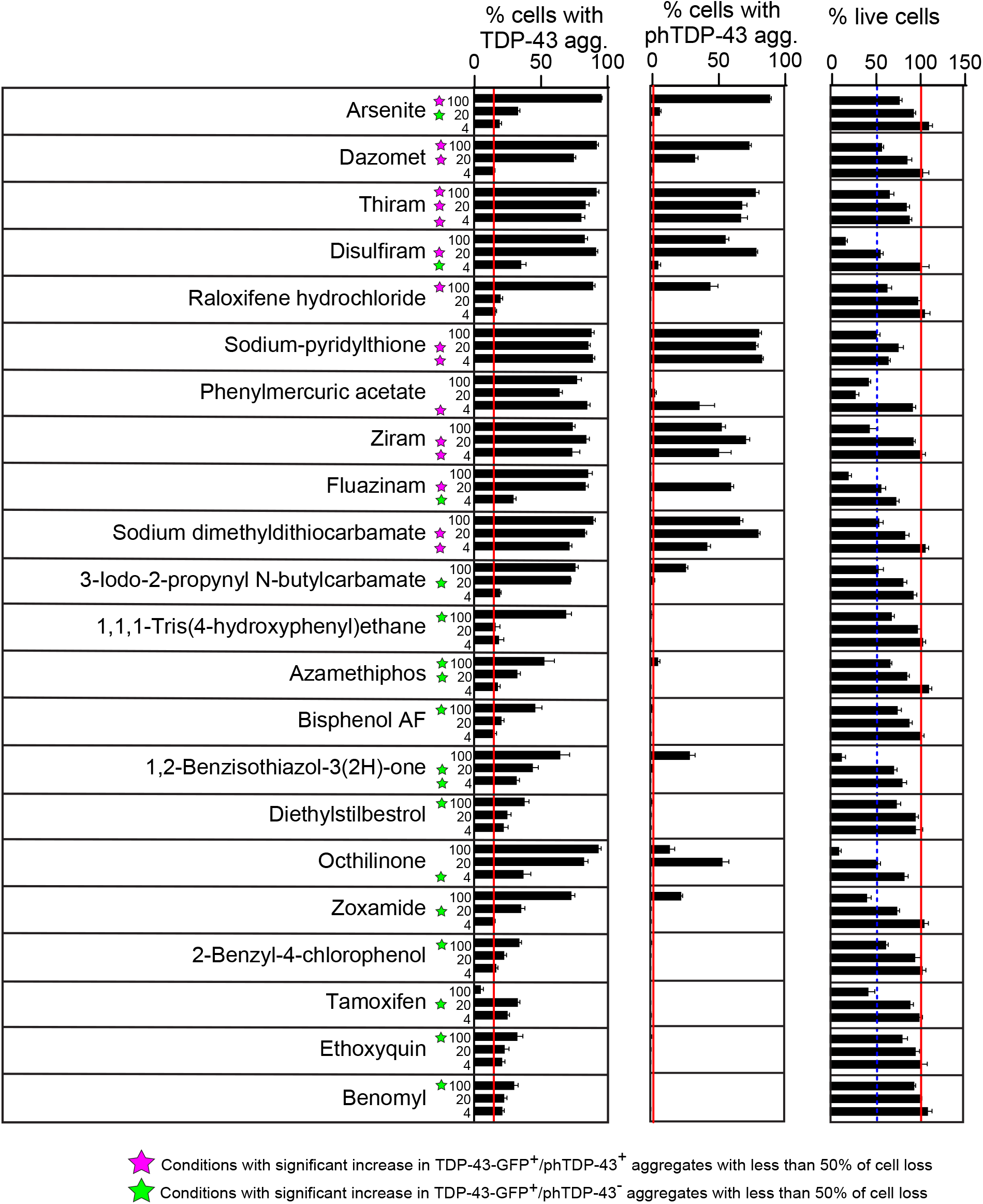
iGFP-TDP43-dNLS validation. Bar chart showing the 21 final hits that induce significant increase in TDP43-GFP aggregation with less than 50% loss in viability and visually validated for the presence of aggregates in at least one dose. All 3 doses are shown. Arsenite is also shown as comparison. Bar charts from left to right show % of cells with TDP43-GFP aggregates, % of cells with phTDP43(S409/10) aggregates and % of number of cells/well relative to DMSO treated cells. In each bar chart the red line indicates average values of the DMSO treated wells. The blue dotted line in the viability bar chart indicates the 50% viability threshold. Conditions that induce a statistically significant increase in just TDP43-GFP aggregates are marked with a green star, conditions that induce a statistically significant increase in phTDP43(S409/10) positive TDP43-GFP aggregates are marked with a magenta star. 2-way ANOVA multiple comparison with Dunnet’s multiple comparison test. n=4 wells/treatment condition, n=12 DMSO control wells. Further statistical information is available in the Statistical Data File. Extended data are available in the Extended Data File

**Supplementary Figure 3.**
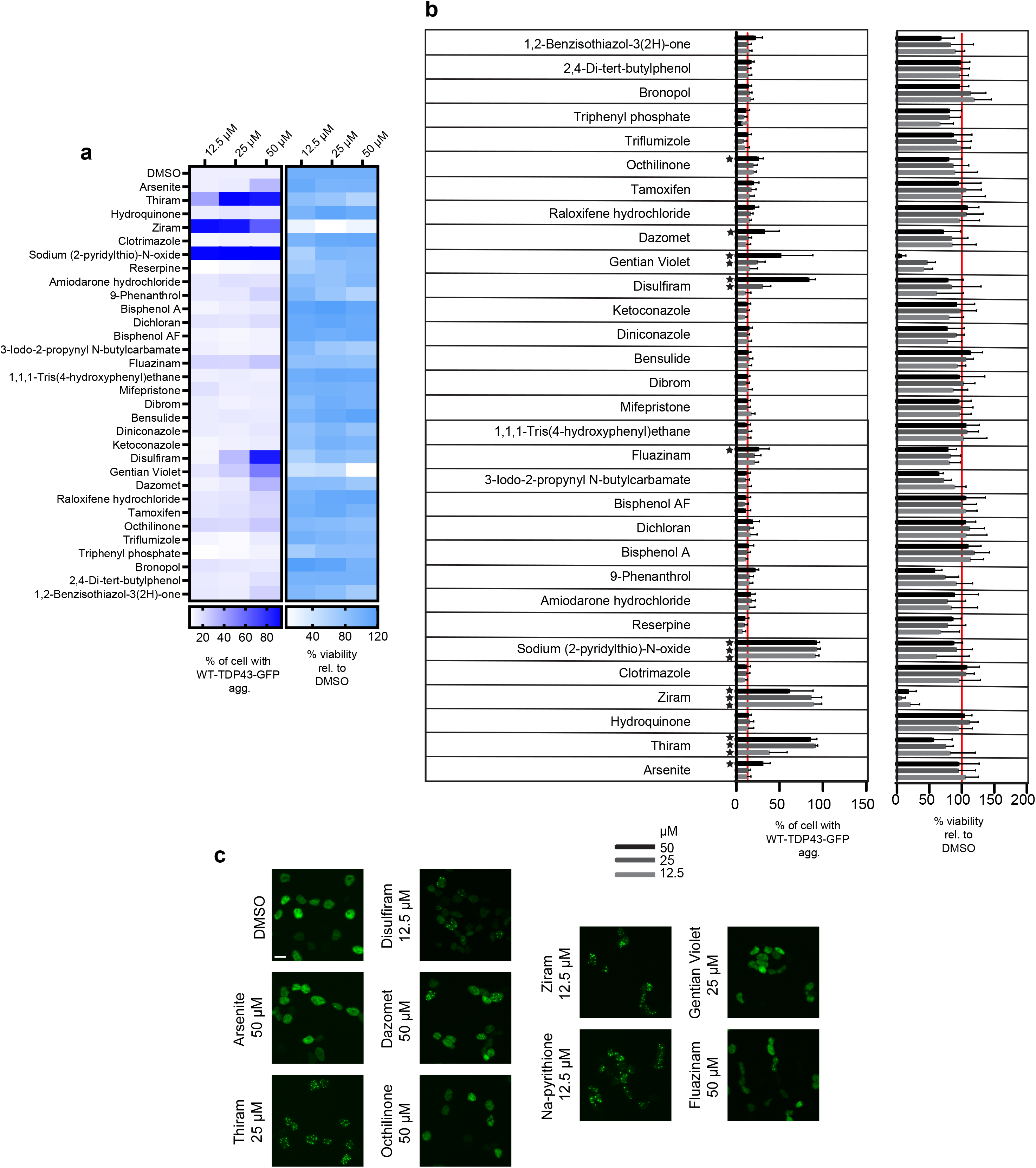
iGFP-TDP43-WT HEKs validation. **a-b** Heat map and bar charts showing changes in the percentage of cells with GFP-TDP43-WT aggregation (left panel) and viability (percentage of cells relative to DMSO wells, right panel) in iGFP-TDP43-WT HEK-293A cells upon treatment with DMSO or 3 concentrations (12.5-25-50 µM, x-axis) of the 30 chemical candidates identified in the screen (listed on y-axis). Arsenite was included as positive control. iGFP-TDP43-WT were treated with 1 µg/ml of doxycycline for 15 hours and exposed to 3 different concentrations of DMSO and chemicals in duplicate wells for 7 hours. Cells were then fixed in 4% PFA and imaged in the GFP/DAPI channels. Red line indicates average value in DMSO control wells. Black stars in bar charts indicate discoveries identified by a 2-way ANOVA test with false discovery rate two stage step-up method of Benjamini, Krieger and Yekutieli correction. n=8 images/treatment condition, n=12 DMSO control images. **c** Representative images showing GFP-TDP43-WT aggregation (green) in iGFP-TDP43-WT HEK-293A cells treated for 7 hours with the 8 chemicals that induce a significant increase in TDP43 aggregation. Scale bar = 25 μm. Further statistical information is available in the Statistical Data File. Extended data are available in the Extended Data File.

**Supplementary Figure 4.**
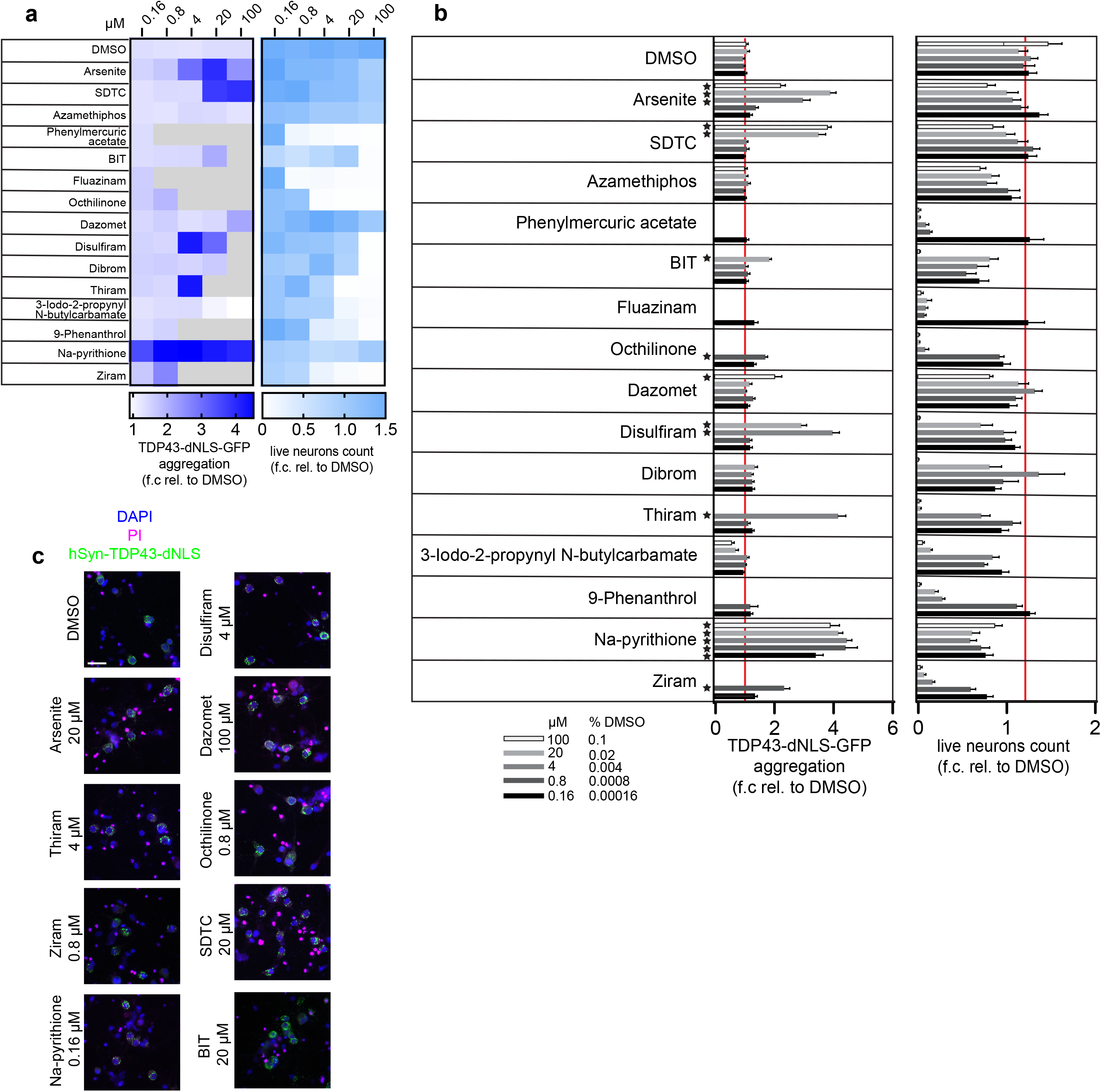
hSyn-TDP43-dNLS-GFP infected neurons validation. **a-b** Heatmap and bar charts showing changes in TDP43-dNLS aggregation (left panel) and viability (right panel) upon treatment with 5 doses (0.16-0.8-4-20-100 µM, x-axis) of DMSO or the best 14 chemical candidates identified in the screen (y-axis). Arsenite was included as positive control. Values are expressed as fold change (f.c.) relative to the average of the lowest DMSO dose (0.00016%). Black stars in bar charts indicate discoveries identified by a 2-way ANOVA test with false discovery rate two stage step-up method of Benjamini, Krieger and Yekutieli correction. n=8 images/condition. **c** Representative images showing TDP43-dNLS-GFP aggregation (green) in hSyn-TDP43-dNLS-GFP infected neurons treated with the 8 chemicals that induce a significant increase in TDP43 aggregation. Lack of PI staining (Magenta) in the nucleus indicates live cells, while DAPI (blue) marks all nuclei. Scale bar = 50 µm. SDTC = sodium dimethyldithiocarbamate, BIT = 2-benzisothi-azol-3(2H)-one. Further statistical information is available in the Statistical Data File. Extended data are available in the Extended Data File.

**Supplementary Figure 5.**
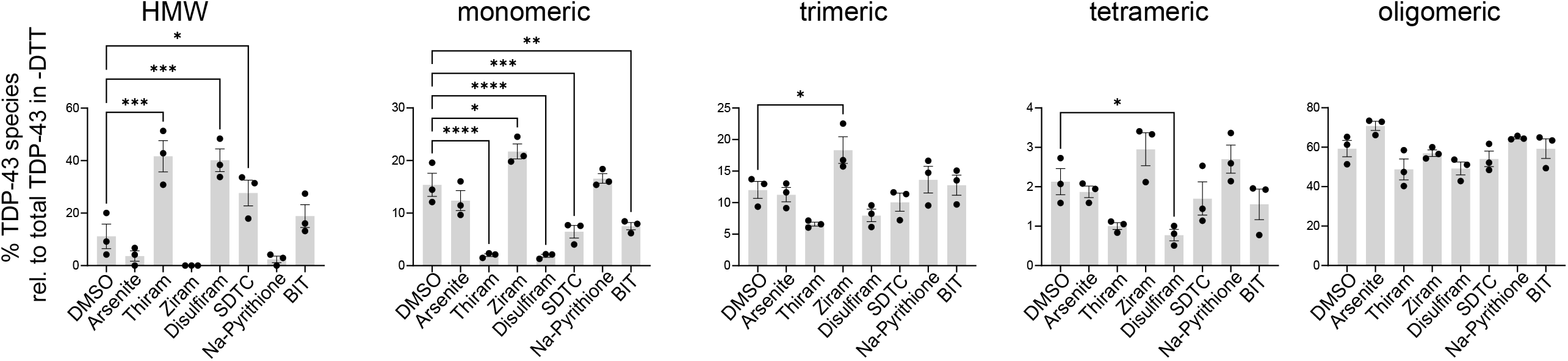
Effect of DTCs and other identified chemicals on rTDP-43 species distribution. Bar-charts show quantification of monomeric (43 kDa), high molecular weight (HMW; >250 kDa), trimeric (∼130 kDa), tetrameric (∼170 kDa), and other oligomeric (present in the smear) TDP-43 species normalized to total amounts of TDP-43 in-DTT blots. Bars represent mean ± SEM; dots represent n=3 biological replicates. One-way ANOVA followed by Dunnet’s multiple comparison test. *p < 0.05, **p < 0.01, ***p < 0.001, ****p < 0.0001. Only significant p values are shown. Further statistical information is available in the Statistical Data File. Extended data are available in the Extended Data File.

**Supplementary Figure 6.**
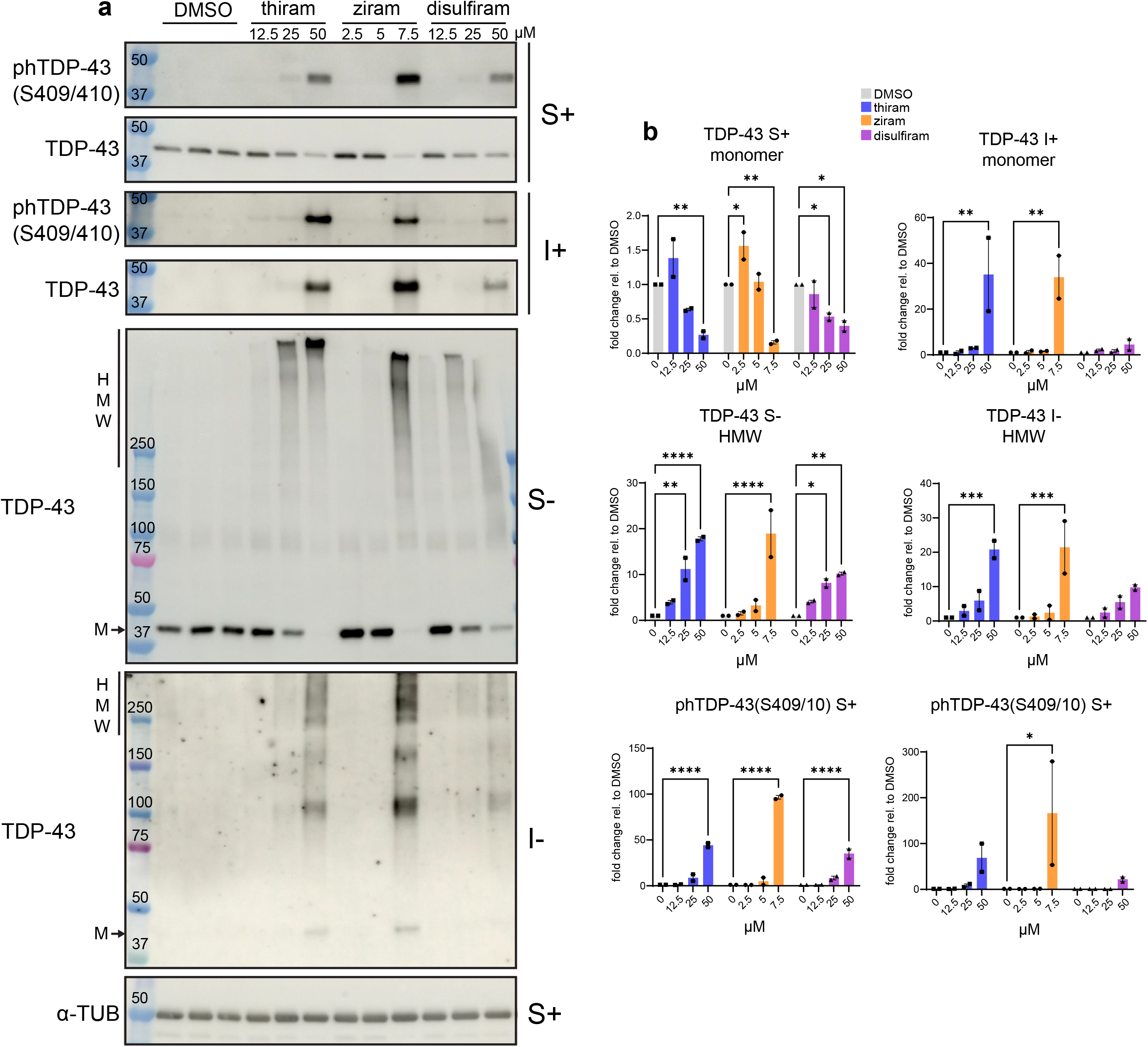
DTCs thiram, ziram and disulfiram induce TDP-43 aggregation in mouse primary neurons. **a** Representative Western blot images of WT mouse cortical neurons treated with DMSO or 3 increasing concentrations of thiram (12.5-25-50 µM), ziram (2.5-5-7.5 µM) or disulfiram (12.5-25-50 µM) for 3 hours. Soluble (S) and insoluble (I) protein fractions were run in presence (+) or absence (-) of DTT. Membranes were blotted with TDP-43, phTDP43(S409/410) and α-TUBULIN (loading control) antibodies. Monomeric (M) and high molecular weight (HMW) TDP-43 species are indicated on the left. **b** Quantification of monomeric (M) or aggregated (HMW) TDP-43 and phTDP43(S409/410) protein levels in soluble and insoluble protein fractions normalized to α-TUBULIN. Bar charts show mean of fold changes in protein levels relative to the average of DMSO samples. n = 2 independent biological replicates. Two-way ANOVA (chemical x concentration) followed by Dunnet’s multiple comparison test. *p < 0.05, **p < 0.01, ***p < 0.001, ****p < 0.0001. Further statistical information is available in the Statistical Data File. Extended data are available in the Extended Data File.

**Supplementary Figure 7.**
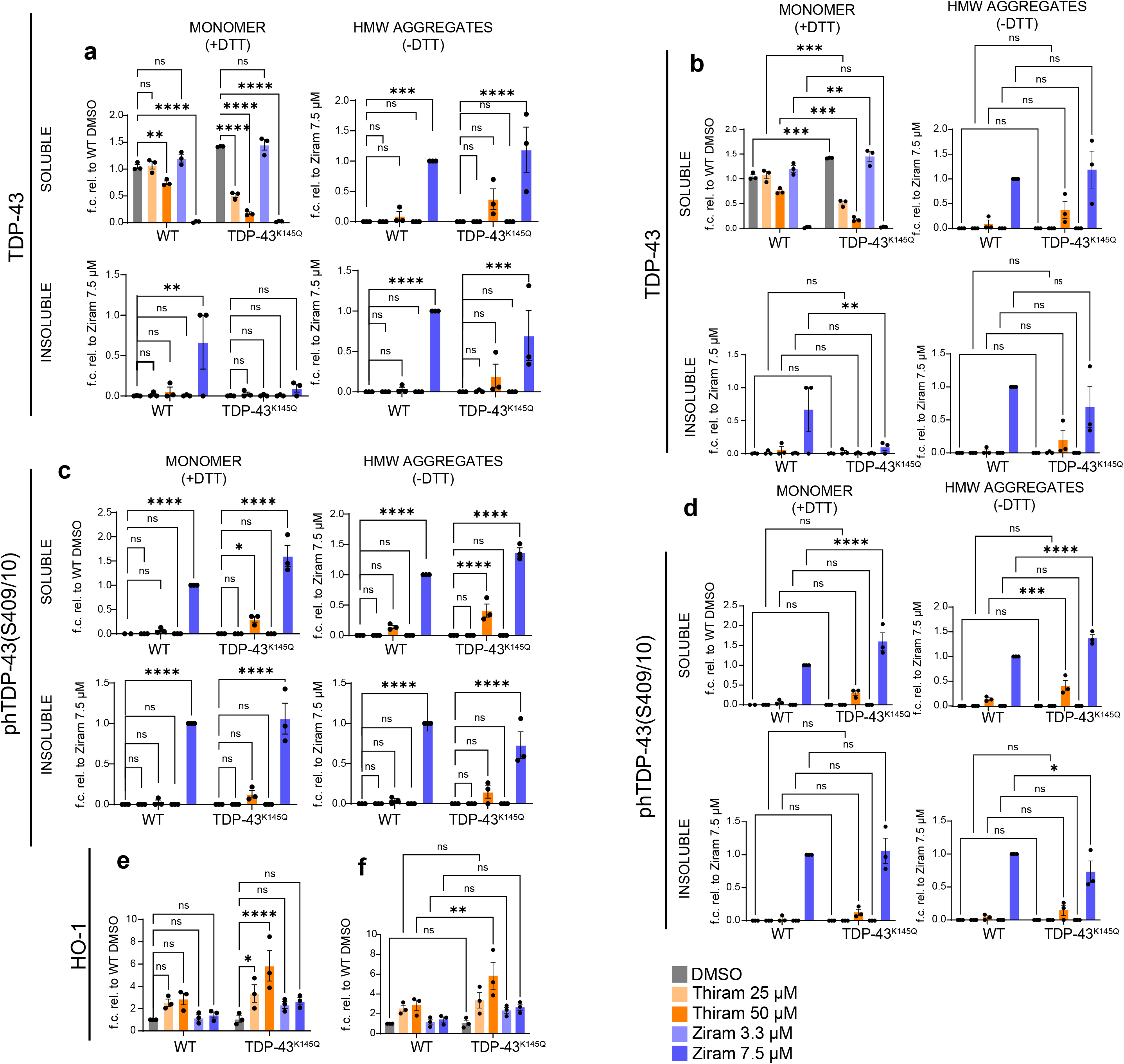
Effect of thiram and ziram treatment on WT and TDP-43 K145Q mouse neurons. Quantifi-cation of monomeric (M) or aggregated (HMW) a-b) TDP-43, c-d), phTDP43(S409/410) and of e-f) HO-1 protein levels in WT and TDP43^K145Q^ mouse cortical neurons treated with DMSO or 2 doses of thiram (25-50 µM) or ziram (3.3-7.5 µM) for 3 hours. Soluble (S) and insoluble (I) protein fractions were run in presence (+) or absence (-) of DTT. Protein amounts are normalized to α-TUBULIN and expressed as fold change relative to the control sample indicated on the y-axis. Two-way ANOVA (genotype x treatment) followed by Dunnet’s multiple comparison test. n = 3 independent biological replicates. Multiple comparisons t-test was performed between the DMSO condition and treatment conditions of the same genotype (a, c, e) or comparing the same treatment condition between two genotypes (b, d, f). ns p>0.05, *p < 0.05, **p < 0.01, ***p < 0.001, ****p < 0.0001. Further statistical information is available in the Statistical Data File. Extended data are available in the Extended Data File.

**Supplementary Figure 8.**
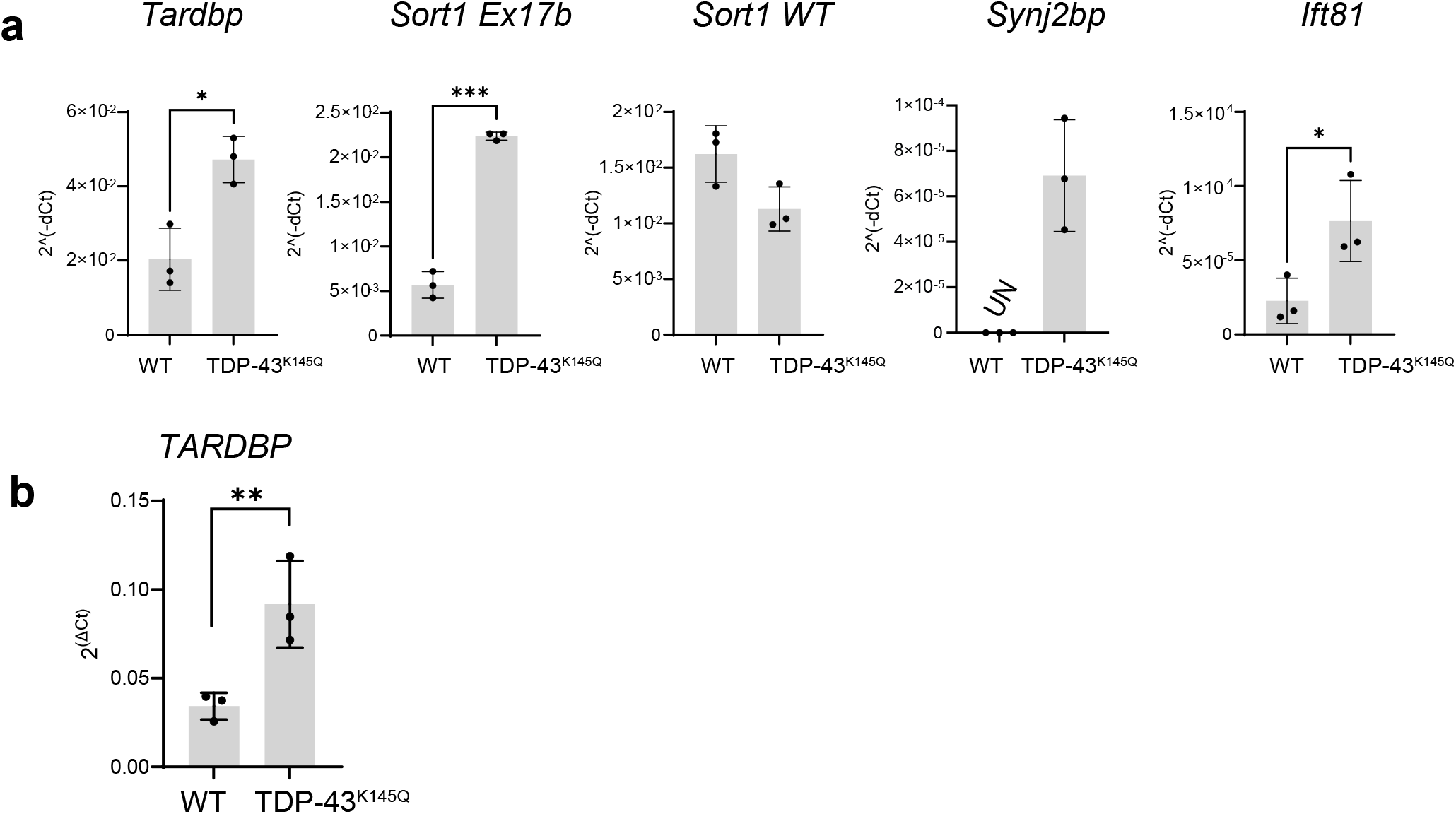
The K145Q mutation induce loss of function of TDP-43 in mouse and hiPSC-derived neurons. qRT-PCR analysis in DMSO treated WT and TDP-43^K145Q^ mouse a) and hiPSC-derived b) neurons showing mRNA levels of the indicated RNA transcripts. mouse mRNA levels were normalized to *Gapdh* housekeeping control. Human mRNA levels were normalized to *RPLP0* housekeeping control. Bars represent mean ± SEM of 2^(-ΔCt)^ values; dots represent n=3 biological replicates. Unpaired t-test on ΔCt values. ns p>0.05, *p < 0.05, **p < 0.01, ***p < 0.001, ****p < 0.0001. *Synj2bp* graph for is provided for visualization purposes only, because non-detectable (UN) levels of Synj2bp cryptic exon-containing transcripts in some or all control TDP-43^WT^ neurons prevented statistical comparisons. Further statistical information is available in the Statistical Data File. Extended data are available in the Extended Data File.

**Supplementary Figure 9.**
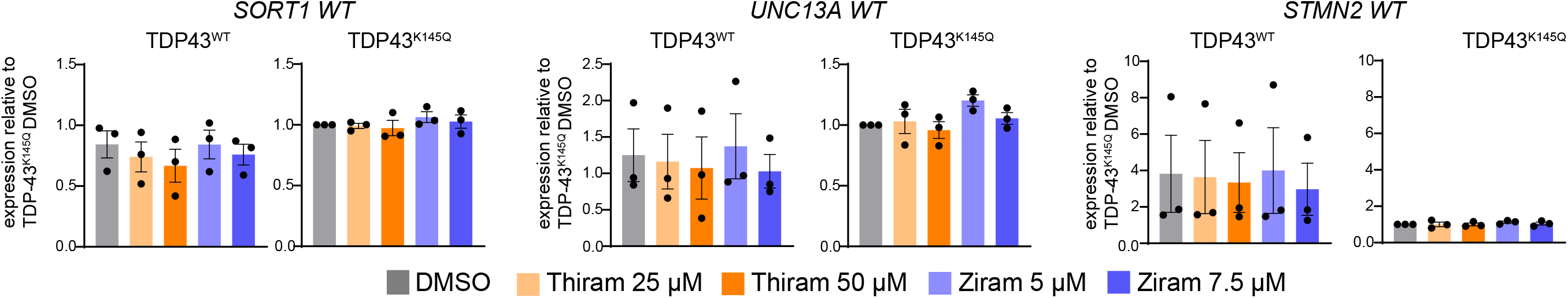
Treatment of hiPSC-derived neurons with thiram and ziram does not affect expression of *SORT1 WT*, *UNC13A WT* and *STMN2 WT* transcripts in hiPSC-derived neurons. qRT-PCR analysis showing amounts of the indicated transcripts in WT and TDP43^K145Q^ hiPSC-derived cortical neurons treated with thiram (25-50 µM) or ziram (5-7.5 µM) for 3 hours. Bars represent mean ± SEM of RNA levels relative to RPLP0 and normalized to TDP43^K145Q^ DMSO sample; dots represent n=3 independent iPSC differentiations and exposures. Friedman test with Dunn’s multiple comparisons test was performed on ΔCt values. *p < 0.05, **p < 0.01, ***p < 0.001, ****p < 0.0001. Only significant p values are shown. Further statistical information is available in the Statistical Data File. Extended data are available in the Extended Data File

**Supplementary Figure 10.**
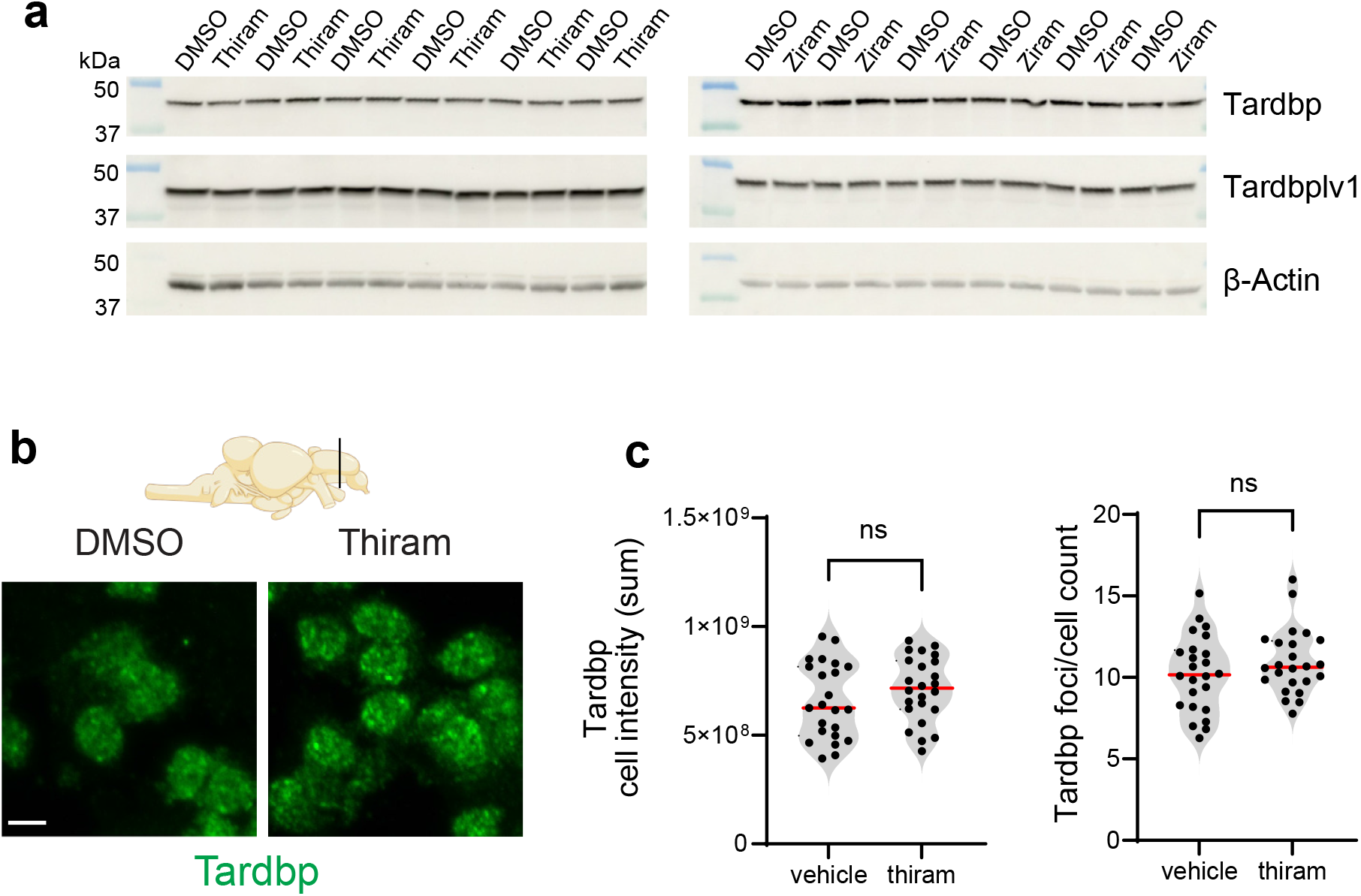
Thiram and ziram exposures did not affect Tardbp and Tardbplv1 protein levels in zebrafish brain. a) Western blot images of Tardbp and Tardpblv1 protein levels in zebrafish brains extracted at 15 days of daily exposures to thiram (0.316 µM) or ziram (1.5 µM). β-Actin was used as loading control. n=6 fish per condition. Representative images b) and quantifications c) of Tarbbp immunofluorescence staining of telencephalic coronal sections of zebrafish exposed for 15 days to thiram (0.316 µM). Scale bar = 5 μm. Dots in violin plots indicate fish mean values. n=24 fish per condition, 5 images per fish. Red bars indicate median values of all fish per condition. Unpaired t-test. ns = p>0.05. Further statistical information is available in the Statistical Data File. Extended data are available in the Extended Data File.

**Supplementary Table 1.**
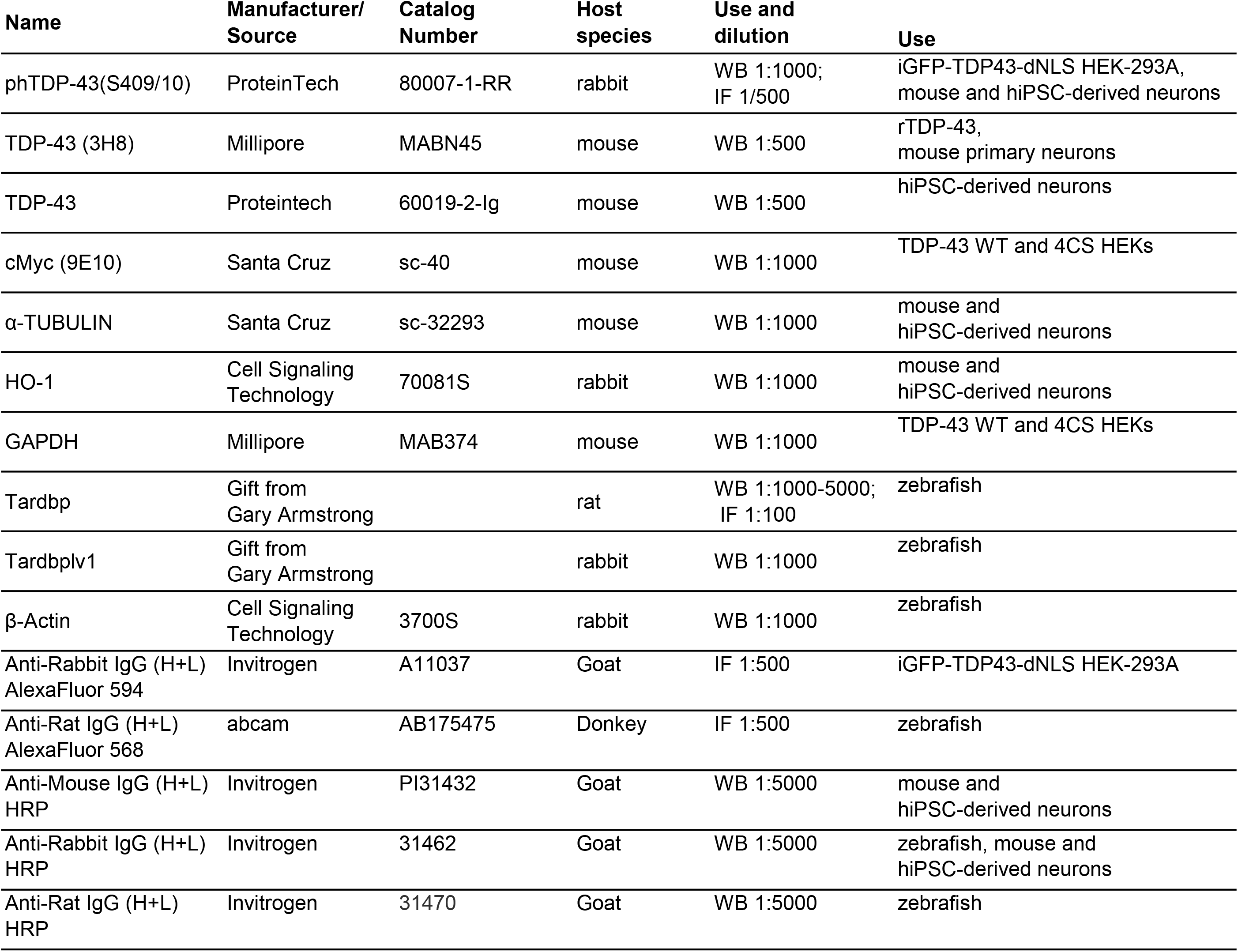
List of antibodies used in this study.

| Name | Manufacturer/<br>Source | Catalog<br>Number | Host<br>species | Use and<br>dilution | Use |
| --- | --- | --- | --- | --- | --- |
| phTDP-43(S409/10) | ProteinTech | 80007-1-RR | rabbit | WB 1:1000;<br>IF 1/500 | iGFP-TDP43-dNLS HEK-293A,<br>mouse and hiPSC-derived neurons |
| TDP-43 (3H8) | Millipore | MABN45 | mouse | WB 1:500 | rTDP-43,<br>mouse primary neurons |
| TDP-43 | Proteintech | 60019-2-Ig | mouse | WB 1:500 | hiPSC-derived neurons |
| cMyc (9E10) | Santa Cruz | sc-40 | mouse | WB 1:1000 | TDP-43 WT and 4CS HEKs |
| α-TUBULIN | Santa Cruz | sc-32293 | mouse | WB 1:1000 | mouse and<br>hiPSC-derived neurons |
| HO-1 | Cell Signaling<br>Technology | 70081S | rabbit | WB 1:1000 | mouse and<br>hiPSC-derived neurons |
| GAPDH | Millipore | MAB374 | mouse | WB 1:1000 | TDP-43 WT and 4CS HEKs |
| Tardbp | Gift from<br>Gary Armstrong |  | rat | WB 1:1000-5000;<br>IF 1:100 | zebrafish |
| Tardbplv1 | Gift from<br>Gary Armstrong |  | rabbit | WB 1:1000 | zebrafish |
| β-Actin | Cell Signaling<br>Technology | 3700S | rabbit | WB 1:1000 | zebrafish |
| Anti-Rabbit IgG (H+L)<br>AlexaFluor 594 | Invitrogen | A11037 | Goat | IF 1:500 | iGFP-TDP43-dNLS HEK-293A |
| Anti-Rat IgG (H+L)<br>AlexaFluor 568 | abcam | AB175475 | Donkey | IF 1:500 | zebrafish |
| Anti-Mouse IgG (H+L)<br>HRP | Invitrogen | PI31432 | Goat | WB 1:5000 | mouse and<br>hiPSC-derived neurons |
| Anti-Rabbit IgG (H+L)<br>HRP | Invitrogen | 31462 | Goat | WB 1:5000 | zebrafish, mouse and<br>hiPSC-derived neurons |
| Anti-Rat IgG (H+L)<br>HRP | Invitrogen | 31470 | Goat | WB 1:5000 | zebrafish |

**Supplementary Table 2.**
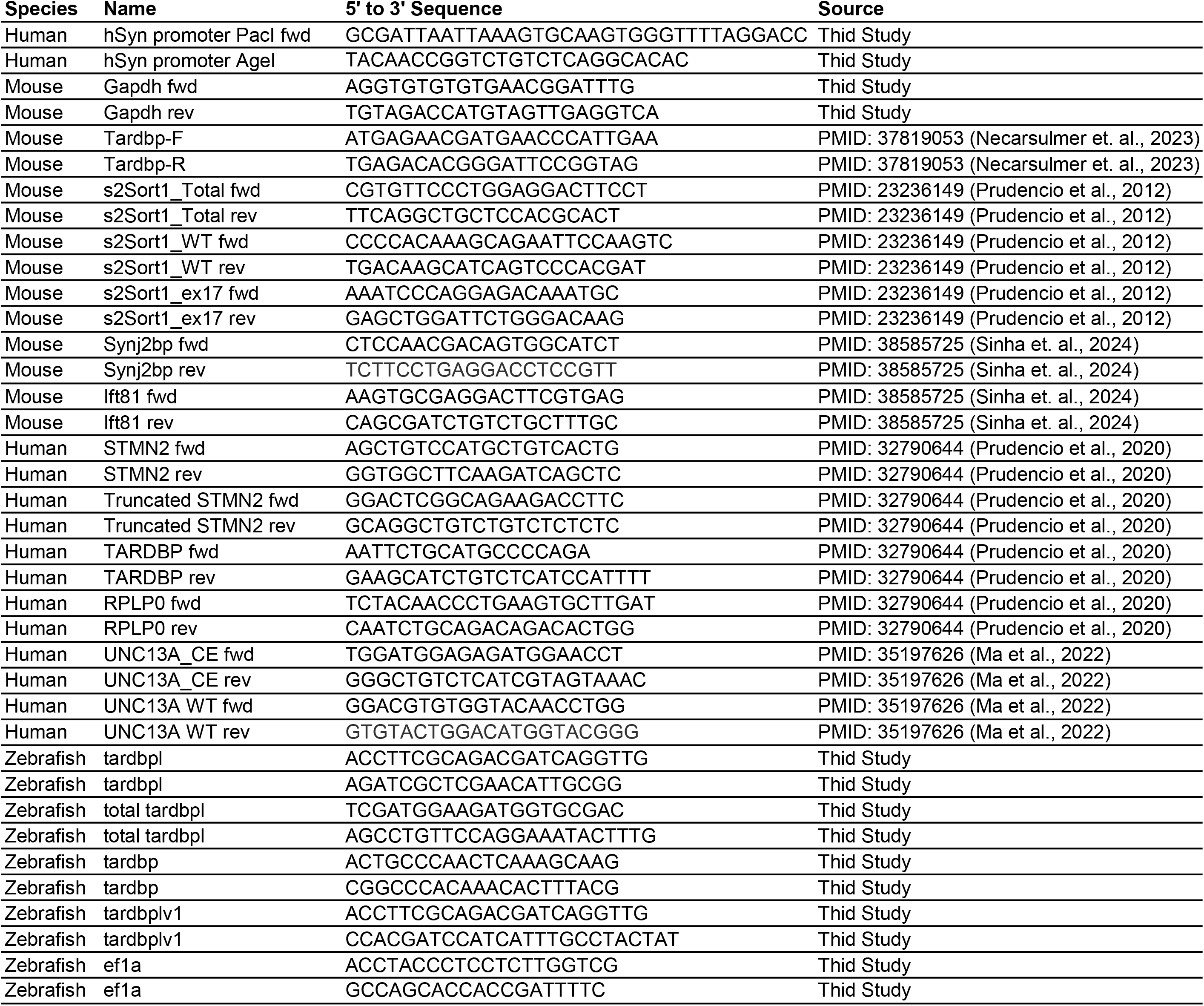
Oligonucleotides used in this study.

